# Structural anatomy of C1 domain interactions with DAG and other agonists

**DOI:** 10.1101/2021.09.03.458901

**Authors:** Sachin S. Katti, Inna V. Krieger, Jihyae Ann, Jeewoo Lee, James C. Sacchettini, Tatyana I. Igumenova

**Affiliations:** Department of Biochemistry and Biophysics, Texas A&M University, TX77840, U.S.A.; College of Pharmacy, Seoul National University, Seoul 08826, Republic of Korea

## Abstract

Diacylglycerol (DAG) is a versatile lipid whose 1,2-*sn*-stereoisomer serves both as second messenger in signal transduction pathways that control vital cellular processes, and as metabolic precursor for downstream signaling lipids such as phosphatidic acid^1,2^. DAG effector proteins compete for available lipid using conserved homology 1 (C1) domains as DAG-sensing modules. Yet, how C1 domains recognize and capture DAG in the complex environment of a biological membrane has remained unresolved for the 40 years since the discovery of Protein Kinase C (PKC)^3^ as the first member of the DAG effector cohort. Herein, we report the first high-resolution crystal structures of a C1 domain (C1B from PKCδ) complexed to DAG and to each of four potent PKC agonists that produce different biological readouts and that command intense therapeutic interest. This structural information details the mechanisms of stereospecific recognition of DAG by the C1 domains, the functional properties of the lipid-binding site, and the identities of the key residues required for the recognition and capture of DAG and exogenous agonists. Moreover, the structures of the five C1 domain complexes provide the high-resolution guides for the design of agents that modulate the activities of DAG effector proteins.

## Introduction

The impressive diversity of DAG signaling output is mediated via its interactions with seven families of effector proteins that execute broad sets of regulatory functions^2^. These include protein phosphorylation (PKCs and PKDs^4,5^); DAG phosphorylation (DGKs^6^); RacGTPase regulation (Chimaerins^7^); Ras guanine nucleotide exchange factor activation (RasGRPs^8^); Cdc42-mediated cytoskeletal reorganization (MRCK^9^); and assembly of scaffolds that potentiate synaptic vesicle fusion and neurotransmitter release (Munc13s^10^). PKCs define a central DAG-sensing node in intracellular phosphoinositide signaling pathways that regulate cell growth, differentiation, apoptosis, and motility^11^. Shortly after their discovery, PKCs were identified as cellular receptors for tumor-promoting phorbol esters^12^ that bind C1 domains in lieu of DAG. These observations, combined with the central roles executed by PKCs in intracellular signaling established their DAG-sensing function as an attractive target for therapeutic intervention, with considerable promise in the treatment of Alzheimer’s disease^13^, HIV/AIDS^14,15^, and cancer^16,17^. However, the structural basis of DAG recognition by the C1 domains remained elusive, and the strategies for therapeutic agent design deployed to date all relied on modeling studies (reviewed in^18^) based on the single available crystal structure of the C1 domain complexed to a ligand that does not activate PKC^19^. Herein, we have overcome these well-documented challenges^20-22^ that have hindered the crystallization of extremely hydrophobic C1-ligand complexes for almost three decades. This advance enabled us to determine high-resolution structures of C1 bound to the endogenous agonist DAG and to each of four exogenous agonists of therapeutic interest. Collectively, our findings: (i) provide a structural rationale for the consensus amino acid sequence of DAG-sensitive C1 domains; (ii) provide insight into the origins of DAG sensitivity; and (iii) reveal how the unique hydrophilic/hydrophobic properties of the ligand-binding groove enable C1 domains to accommodate chemically diverse ligands.

### Structure of the C1Bδ-DAG complex

C1 domains form complexes with their ligands in the membrane environment, and dodecylphosphocholine (DPC) is an effective membrane mimic that faithfully reproduces the functional properties of C1 domains with respect to ligand interactions^23,24^. For structural studies, we formed the complex between the C1B domain of PKCδ (C1Bδ) and a synthetic DAG analog (dioctanoyl-*sn*-1,2-glycerol) in the presence of DPC micelles. The structure of the C1Bδ-DAG complex was refined to 1.75 Å with R_work_=0.214 and R_free_=0.246 (**SI Table 1**). The H3 space group unit cell (89 × 89 × 219 Å, **Fig. 1a**) contains a total of 72 protein chains contributed by nine asymmetric units (AUs) with eight C1Bδ molecules per AU (**Fig. 1b**). The crystal lattice organization is unusual as it highlights two distinct lipid-detergent micelles: micelle 1 composed of 12 DAG and 12 DPC ordered molecules, and micelle 2 composed of 18 DAG and 6 DPC ordered molecules (**Fig. 1c**), in addition to fully or partially disordered lipids. We speculate that the micelles help nucleate the crystallization, as all of the protein subunits are arranged with their lipid sensing loops directly binding to DAG within micelles (**Ext. Figure 1a,b**). Each C1Bδ protein chain has a DAG molecule captured within a groove formed by its membrane-binding loop regions (**Fig. 1b** and **Ext. Fig. 1c,d**). The well-defined glycerol ester moieties of this tightly bound ‘intra-loop’ DAG refined with B-factors (18-26 Å) comparable to those of the surrounding protein residues (17-23 Å). In addition, each AU contains two less ordered peripheral DAG and six DPC molecules associated with the amphiphilic protein surface (**Ext. Fig. 1e,f**).

**Fig.1.**
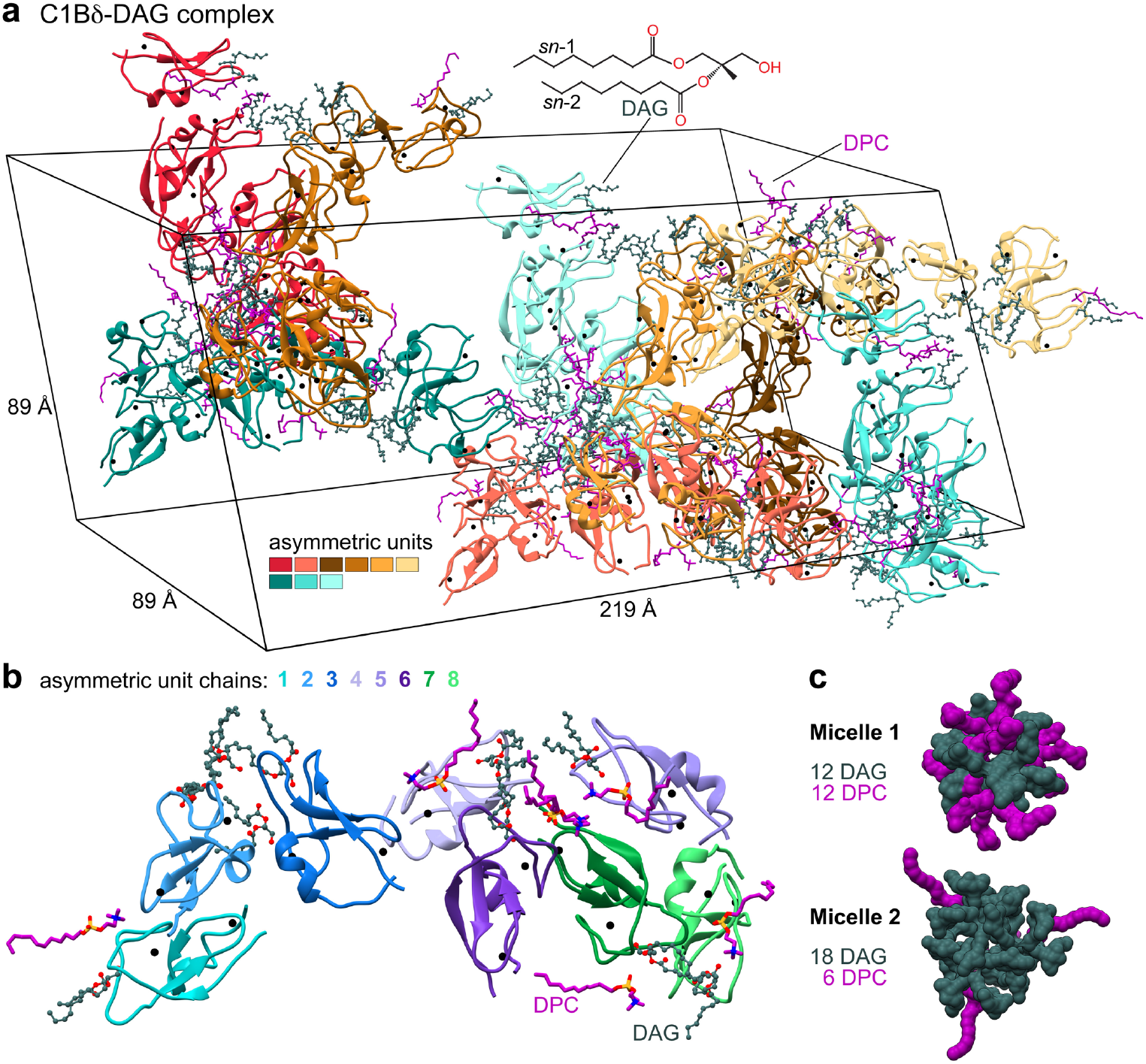
Arrangement of protein chains, lipid, and detergent molecules in the unit cell and the asymmetric unit of the C1Bδ-DAG complex crystal (PDB ID: 7L92). **a**, The unit cell contains 72 DAG-complexed C1Bδ chains and 18/54 DAG/DPC molecules that peripherally associate with the protein surface. Structural Zn^2+^ ions of C1Bδ are shown as black spheres. **b**, The asymmetric unit comprises 8 C1Bδ protein chains with 8 DAG molecules captured within a well-defined groove, and 2/6 peripheral DAG/DPC molecules. **c**, Space-filling representation of the two distinct DAG/DPC micelles.

There is little variability among DAG-complexed C1Bδ chains within the asymmetric unit, as evidenced by low pairwise backbone r.m.s.d values of 0.3-0.6 Å (**Fig. 2a**). The most variable region is located between helix α1 and the C-terminal Cysteine residue that coordinates a structural Zn^2+^ ion. According to solution NMR, this region undergoes conformational exchange on the μs-timescale^23,25^. Apo C1Bδ was also crystallized under conditions similar to those for the C1Bδ-DAG complex. The apo structure superimposes well onto the structures of DAG-complexed C1Bδ, with the notable exception of the Trp252 sidechain. In the DAG complex, this sidechain is oriented towards the DAG tethered to the membrane-binding region, whereas in the apo C1Bδ it is oriented away from that region (**Fig. 2a**). This was a satisfying result as Trp252 is associated with the “DAG-toggling” behavior of the C1 domains, wherein a conservative Trp→Tyr substitution in novel (or Ca^2+^-insensitive) and Tyr→Trp substitution in conventional (or Ca^2+^-activated) PKC isoforms significantly modulates apparent affinity for DAG^23,24,26,27^.

**Fig.2.**
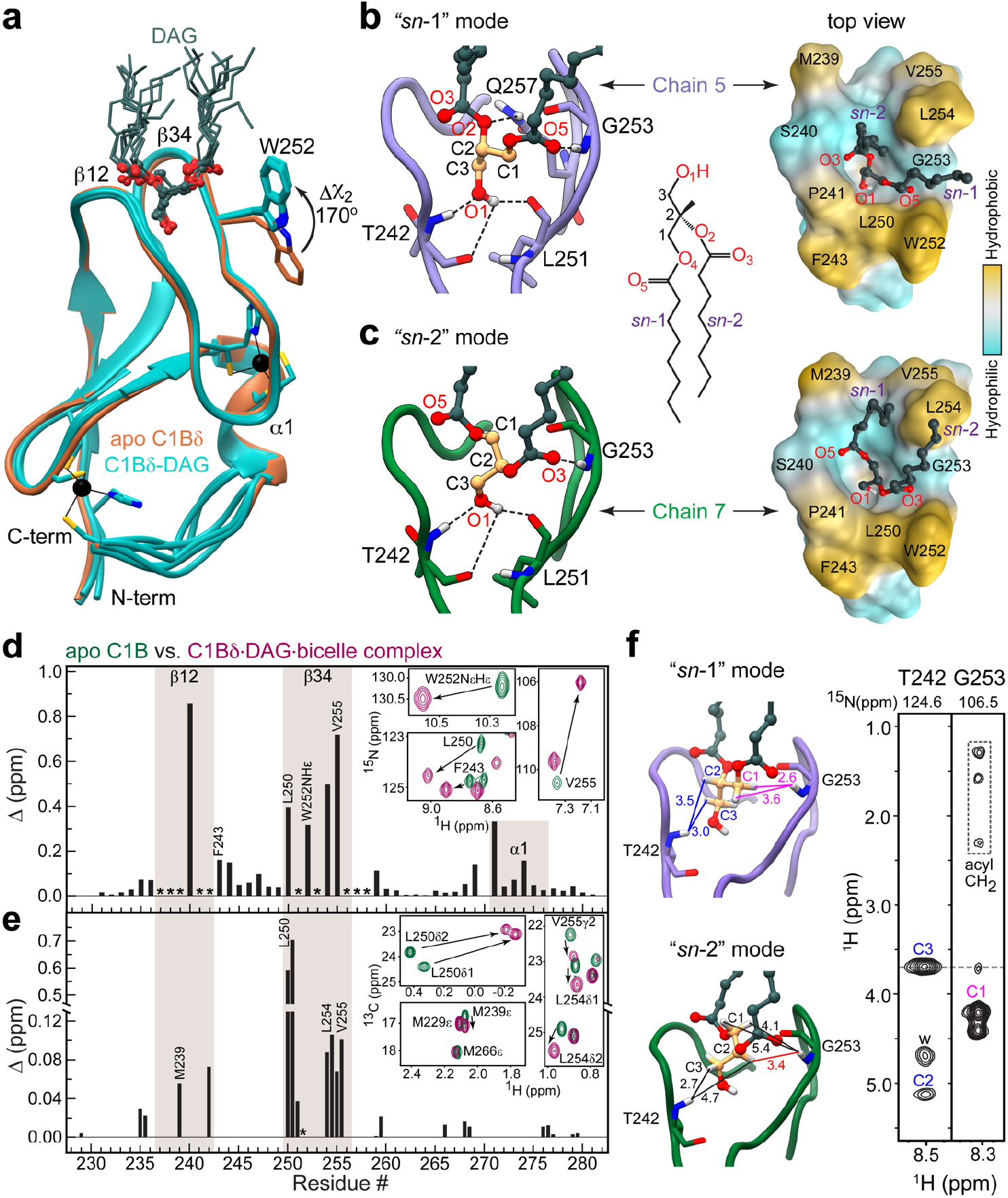
Stereospecificity of DAG binding by C1Bδ. **a**, Backbone superposition of 8 DAG-complexed C1Bδ chains of the AU (cyan, PDB ID: 7L92) onto the structure of apo C1Bδ (sienna, PDB ID: 7KND). The sidechain of Trp252 reorients towards the tips of membrane-binding β12 and β34 loops upon DAG binding. DAG adopts one of the two distinct binding modes: “*sn*-1” (**b**) or “*sn*-2” (**c**). The formation of the C1Bδ-DAG complex in bicelles is reported by the chemical shift perturbations (CSPs) of the amide ^15^NH (**d**) and methyl ^13^CH_3_ (**e**) groups of C1Bδ. Asterisks denote residues whose resonances are broadened by chemical exchange in the apo state. The insets show the response of individual residues to DAG binding through the expansions of the ^15^N-^1^H and ^13^C-^1^H HSQC spectral overlays of apo and DAG-complexed C1Bδ. **f**, ^1^H-^1^H_N_ Thr242 and Gly253 strips from the 3D ^15^N-edited NOESY-TROSY spectrum of the C1Bδ-DAG-bicelle complex. The protein-to-DAG NOE pattern is consistent with the distances observed in the “*sn-*1” mode (Chain 5, light purple) but not the “*sn*-2” mode (Chain 7, green). All distances are in Å and color-coded in the “*sn*-1” complex to match the labels in the spectrum; “w” denotes water protons. The medium-range NOE that would be characteristic of the “*sn*-2” complex is shown in red.

### Stereospecificity of DAG binding by C1Bδ

Another significant feature of the C1Bδ-DAG structure is that it reveals the mechanism for stereospecific binding of *sn*-1,2-diacylglycerol by C1 domains. DAG binds in the groove formed by the protein loops, β12 and β34 (**Fig. 2a**) and can adopt two distinct binding modes: “*sn*-1” and “*sn*-2” (**Figs. 2b,c**)^28^. The “*sn*-1” mode is predominant in the crystal as it is represented in six of the eight C1Bδ chains (**Ext. Fig. 2a**). We note that the DAG/DPC micelle 1 exclusively supports the “*sn*-1” binding mode, while micelle 2 supports both, “*sn*-1” and “*sn*-2” (**Ext. Fig. 1b**).

In both modes, the DAG glycerol and ester moieties are anchored to the C1Bδ binding groove by four hydrogen bonds. Three are contributed by the C3-OH hydroxyl group that serves as the donor for the carbonyl oxygens of Thr242 and Leu251, and as the acceptor for the amide hydrogen of Thr242. The fourth bond involves the amide hydrogen of Gly253 and it is this bond that defines the binding mode. In the “*sn*-1” position, the acceptor is the carbonyl oxygen O5 of the *sn*-1 ester group, whereas in the “*sn*-2” position the acceptor is the carbonyl oxygen O3 of the *sn*-2 ester group (**Fig. 2b,c**).

A particularly important feature of the “*sn*-1” binding mode is involvement of the alkoxy oxygen O2 of the *sn*-2 DAG chain in the hydrogen bond with the amide protons of the Gln257 sidechain (**Fig. 2b, Ext. Fig. 2b**). Gln257 is part of the strictly conserved “QG” motif in all DAG-sensitive C1 domains^29^ that is essential for agonist binding^21,23,30,31^, and whose sub-millisecond dynamics in apo C1 domains correlates with DAG binding affinity^23^. The Gln257 sidechain also “stitches” the β12 and β34 loops together, forming hydrogen bonds with Tyr238 of loop β12 and Gly253 of loop β34 (**Ext. Fig. 2b**). The simultaneous involvement of Gln257 in both DAG and intra-protein stabilizing interactions explains its essential role in the formation of the C1Bδ-DAG complex.

### C1Bδ binds DAG in the “*sn*-1” mode in solution

To determine the predominant DAG binding mode in solution, we conducted solution NMR experiments on C1Bδ-DAG assembled in isotropically tumbling bicelles. Complex formation is evident from the chemical shift perturbations of the C1Bδ backbone NH and the Trp252 NHε groups (**Fig. 2d** and **Ext. Fig. 3a**), and of the methyl groups of hydrophobic residues residing in the loop regions (**Fig. 2e** and **Ext. Fig. 3b**). Also evident is rigidification of the loops upon DAG binding as manifested by the appearance of backbone NH cross-peaks broadened in the apo state due to their intermediate-timescale loop dynamics (**Ext. Fig. 3a**). 3D ^15^N-edited [^1^H,^1^H] NOESY experiments were performed where C1Bδ was extensively deuterated at the non-exchangeable sites to suppress intra-protein NOEs. The two DAG binding modes observed in the crystal structure will produce drastically different protein-to-ligand NOE patterns. The signatures of the “*sn*-1” mode are predicted to be short-to-medium range NOEs between ^1^H_N_(Gly253) and ^1^H_CH2_(C1), and a long-range NOE between ^1^H_N_(Gly253) and ^1^H_CH2_(C3). This is precisely the pattern observed experimentally in the Gly253 strip (**Fig. 2f**). The medium-range NOE between ^1^H_N_(Gly253) and ^1^H_CH_(C2) that would signify the “*sn*-2” mode (shown in red in the “*sn*-2” complex, **Fig. 2f**) is not detected. Further confirmation of the “*sn*-1” DAG binding mode is provided by the Thr242 strip whose ^1^H_N_ shows a characteristic medium-range NOE to ^1^H_CH_(C2) and a short-range NOE to ^1^H_CH2_(C3). Thus, both the C1Bδ-DAG structure and solution NMR experiments report “*sn*-1” as the dominant mode of DAG binding to C1Bδ. Moreover, also consistent with the C1Bδ-DAG structure, several C1Bδ loop residues (including Gly253) show NOEs to the methylenes of acyl chains of either DAG or bicelle lipids (**Ext. Fig. 3c,d**).

### Roles of C1Bδ loops in lipid binding

Inspection of C1Bδ loop regions in the C1Bδ-DAG structure reveals how exquisitely they are tuned to the chemical properties of diacylglycerol and surrounding lipids. The eleven C1Bδ residues involved in DAG interactions can be grouped into three tiers that progressively increase the hydrophobicity of the ligand environment – as illustrated using the deconstructed binding groove of the representative “*sn*-1” complex (**Fig. 3a**). The atoms from four “tier 1” residues, namely the backbone O and NH groups of Tyr238, Thr242, and Leu251, along with the sidechain of Gln257, define the polar surface of the groove floor that accommodates the C3-OH hydroxyl group of DAG. The hydrophobic sidechains of tier 1 residues point towards the core of the protein and away from the groove. The five “tier 2” residues accommodate the glycerol backbone and ester groups of DAG by creating an amphiphilic environment. Met239 and Ser240 backbone O and N atoms provide a polar environment for the O3 oxygen of DAG, while their sidechains face “outward” to potentially engage in lipid interactions. Pro241, a strictly conserved residue in DAG-sensitive C1 domains, makes non-polar contacts with the C2 carbon of DAG, and its Cδ/Cα are positioned sufficiently close to the DAG oxygens to form C-H…O interactions (**Ext. Fig. 4c**). On the opposite side of the groove, the polar N-H group of Gly253 hydrogen bonds to the O5 oxygen, while the hydrophobic Leu250 sidechain engages in non-polar contacts with the C1 carbon of DAG.

**Fig.3.**
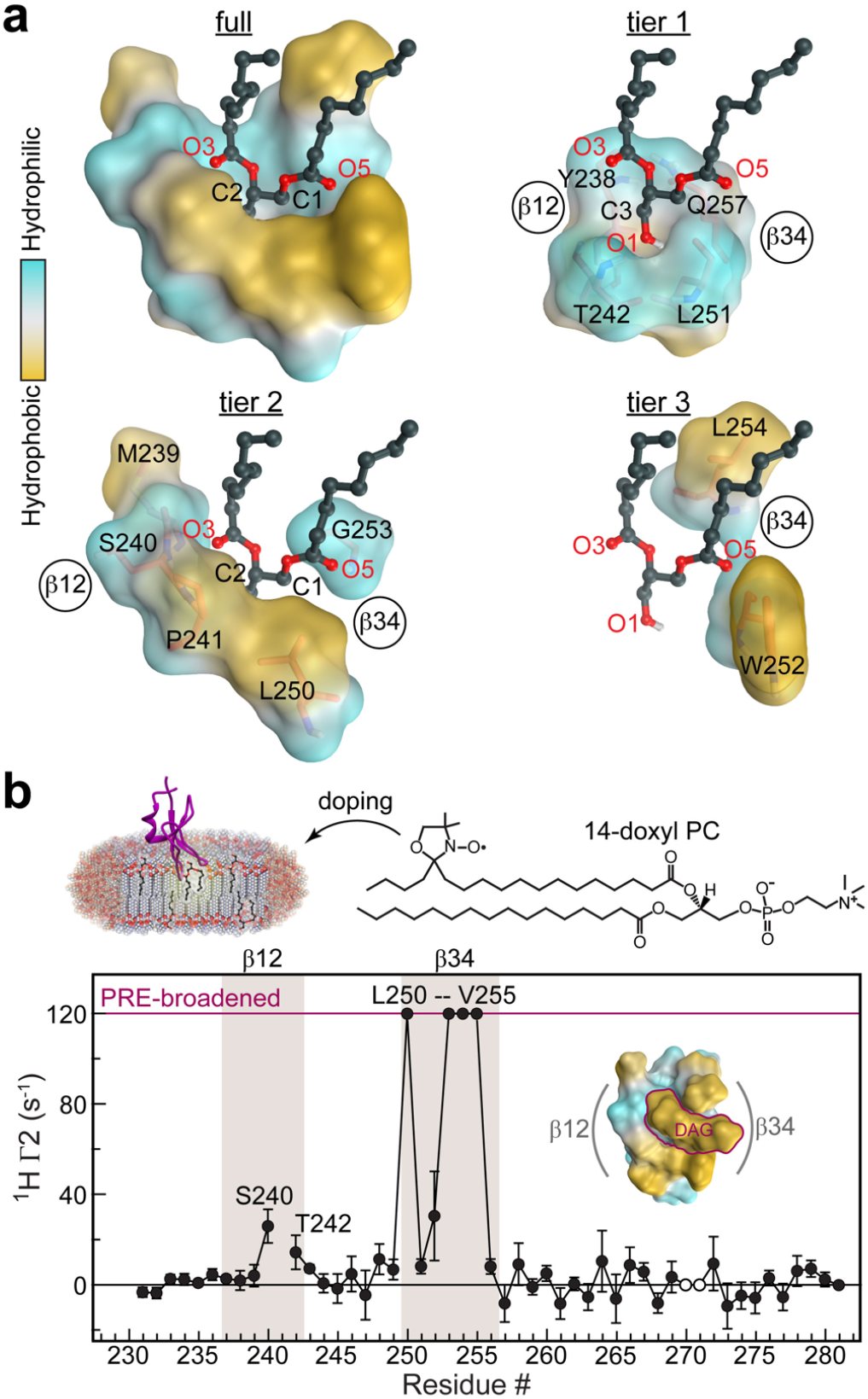
Roles of C1Bδ loops in lipid binding. **a**, Polar backbone atoms and hydrophobic sidechains of DAG-interacting C1Bδ residues create a binding site whose properties are tailored to capture the amphiphilic DAG molecule. This is illustrated through the deconstruction of the “*sn-1*” binding mode into three tiers that accommodate the glycerol backbone (tier 1), the *sn*-1/2 ester groups (tier 2), and the acyl chain methylenes (tier 3). **b**, Residue-specific lipid-to-protein PRE values of the amide protons, ^1^H r_2_, indicate that loop β34 is inserted deeper into the membrane than β12. The PRE value for Trp252 is that for the indole NHε group. Cross-peaks broadened beyond detection in paramagnetic bicelles are assigned an arbitrary value of 120 s^-1^. His270 and Lys271 cross-peaks are exchange-broadened and therefore unsuitable for quantitative analysis (open circles). The inset shows the top view of the “*sn-1*” mode C1Bδ-DAG complex color-coded according to the hydrophobicity.

While tier 1 and 2 residues are contributed by both loops, tier 3 residues all reside on loop β34 (**Fig. 3a**). Trp252 and Leu254 sidechains make nonpolar contacts with the methylenes of the DAG *sn*-1 and *sn*-2 acyl chains (**Ext. Fig. 4a**) that protrude through the depression formed by Gly253 (**Fig. 2b, 3a**). These interactions orient the DAG acyl chains in a position to complete the hydrophobic rim of the C1Bδ domain formed by loop β34 residues Leu250, Trp252, Leu254, Val255, and loop β12 residues Met239 and Pro241. This is illustrated in top views of the “*sn*-1” and “*sn*-2” C1Bδ-DAG complexes (right panels of **Fig. 2b,c**). Positioning the acyl chains in close proximity to Trp252 and Leu254 sidechains creates a continuous hydrophobic surface tailored for C1Bδ interactions with surrounding lipids. In this context, the significance of Trp252 sidechain reorientation upon DAG binding (**Fig. 2a**) becomes clear, as this conformation ensures the continuity of the hydrophobic surface. A direct manifestation of the lipophilicity of the DAG-bound C1Bδ surface is the peripheral association of DAG and detergents observed in all crystal structures of the complexes (**Ext. Fig. 5, 6**, and **10**; **Extended discussion 2**).

To directly identify the regions of the C1Bδ-DAG that insert into the bilayer and to independently validate the essential role of the β34 loop in membrane partitioning, paramagnetic relaxation enhancement (PRE) experiments were performed using a paramagnetic lipid (14-doxyl PC) incorporated into host bicelles. PREs arise due to spatial proximity of the unpaired electron of the lipid probe to protein nuclear spins, and manifest themselves as extensive line broadening in the NMR spectra. PRE data for the C1Bδ-DAG amide hydrogens report that, while both loops undergo bilayer insertion, loop β34 penetrates deeper into the bilayer than does loop β12 (**Fig. 3b**). The PRE data are entirely consistent with the hydrophobicity patterns observed in the crystal structure (**Fig. 3b, inset**), and project that C1Bδ assumes a tilted position relative to the membrane normal upon DAG binding.

Of note, the deconstructed binding groove of the “*sn*-2” complex shows a similar three-tiered arrangement of hydrophilic and hydrophobic features equally suited to accommodate DAG in the “*sn*-2” orientation (**Ext. Fig. 4b,d**). However, because of the differences in the hydrogen-bonding patterns (left panels of **Figs. 2b,c**) DAG position is shallow compared to the “*sn*-1” mode, where the DAG C1 carbon resides deeper in the pocket by ∼1.5 Å. While our data support the “*sn*-1” as the dominant DAG interaction mode, the presence of the “*sn*-2” mode in the crystal structure (**Ext. Fig. 1b, 2a**) suggests that it too is sampled transiently during the DAG capture step.

### C1Bδ complexes with exogenous PKC agonists

The DAG-sensing function has been an active target for pharmaceutical modulation of PKC activity. To determine the structural basis of how C1 domains mediate the response of DAG effector proteins to potent exogenous agonists, we selected four such agonists that evoke distinct cellular responses. Phorbol 12,13-dibutyrate (PDBu) is one of the potent tumor-promoting phorbol esters widely used to generate carcinogenesis models through PKC dysregulation^32^. Prostratin, a non-tumorigenic phorbol ester, is a pre-clinical candidate for inducing latency reversal in HIV-1 infection^33,34^. Both PDBu and Prostratin share a tetracyclic tigliane skeleton of 5-7-6-3 membered rings (**Fig. 4a**). Ingenol 3-angelate (I3A) is a clinically approved agent for the topical treatment of actinic keratosis (Picato®) with a phorbol-related 5-6-6-3 fused ring structure^35^. AJH-836 is a high-affinity synthetic DAG lactone with considerable promise as an isoenzyme-specific PKC agonist (**Extended discussion 3**)^36^. In the membrane-mimicking lipid bicelle environment, all four ligands bind to the loop region of C1Bδ – as evidenced by the chemical shift perturbations and rigidification of the corresponding residues (**Ext. Fig. 7**). Crystals of each C1Bδ-ligand complex formed in the presence of 1,2-diheptanoyl-sn-glycero-3-phosphocholine (DHPC) and all yielded high-resolution structures (1.1 to 1.8 Å, **SI Tables 1-2**) with well-defined electron densities of ligands (**Ext. Fig. 8a**).

**Fig.4.**
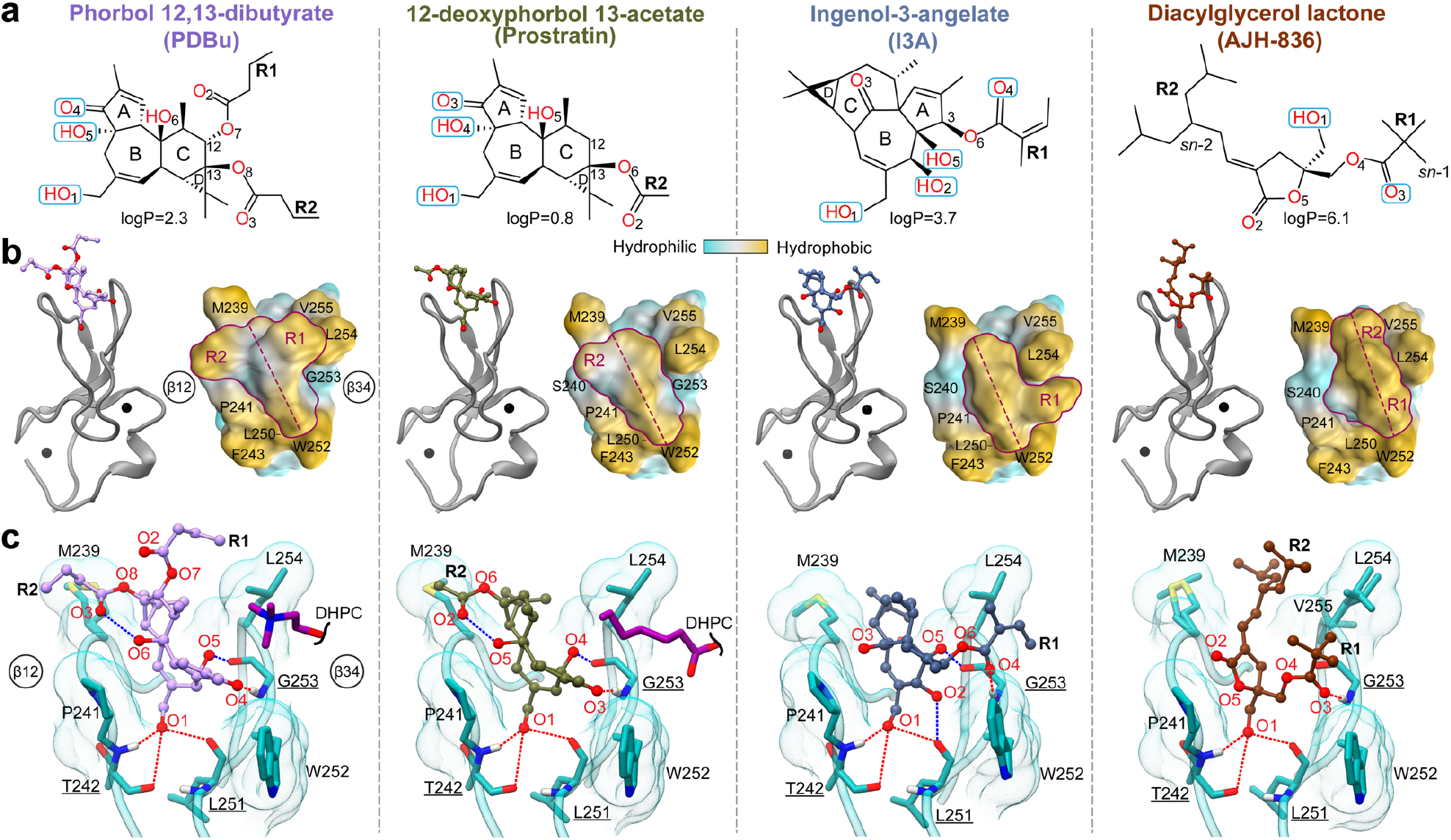
Structures of the C1Bδ-ligand complexes reveal the interaction modes of PKC agonists. **a**, Chemical structures and polar groups involved in hydrogen-bonding interactions with C1Bδ of PKC agonists. The numbering of oxygen atoms follows the ALATIS system. **b**, 3D structures of the complexes (PDB IDs from left to right: 7KNJ, 7LCB, 7KO6, and 7LF3) showing the ligand placement in the binding groove. The shape of the ligands’ hydrophobic cap, viewed from the top of the loop region, is outlined in maroon. The hydrophobic ridge that traverses the groove is marked with the maroon dashed line. **c**, Ligand interactions with Thr242, Leu251, and Gly253 (underlined) that recapitulate the DAG hydrogen-bonding pattern are shown with red dashed lines. Blue dashed lines show ligand-specific hydrogen bonds, including the intra-ligand ones in PDBu and Prostratin. The depression created in loop β34 by Gly253 in the PDBu and Prostratin complexes accommodates DHPC molecules in the crystal.

These structures reveal that all four ligands bind to the same C1Bδ site as DAG (**Fig. 4b**). However, none of them form a hydrogen bond with the Gln257 sidechain that is observed in the DAG complex structure (**Fig. 2b**). The fused ring structures of PDBu, Prostratin, and I3A intercalate between loops β12 and β34. The methyl groups attached to rings A, C, and D, and the apical regions of rings A and C (Prostratin and I3A only) protrude outwards from the groove (**Ext. Fig. 8b**). These groups collectively form a hydrophobic ridge that traverses the groove diagonally from Met239 of loop β12 to Trp252 of loop β34 (**Fig. 4b**). The AJH-836 lactone ring also intercalates between the loops and is fully sequestered.

The arrangement and identity of the R groups provide unique shape of the hydrophobic “cap” over the loop region (**Fig. 4b**, surface representation). In the PDBu complex, the butyryl R1 and R2 groups are arranged in a T-shape relative to the ridge. Prostratin, having only a single acetyl R2 group, forms a significantly smaller hydrophobic cap. Neither ligand covers the depression in the β34 loop formed by Gly253 – thereby creating an opportunistic interaction site for DHPC molecules in the crystal (**Fig. 4c**). In contrast, bound I3A and AJH-836 form a contiguous hydrophobic surface with loop β34, while leaving loop β12 exposed and available for potential interactions with lipids (**Fig. 4b**). The angelyl and pivaloyl R1 groups of I3A and AJH-836, respectively, occupy the depression created by Gly253 and engage in hydrophobic contacts with the sidechains of the bracketing residues Leu254 and Trp252 (**Ext. Fig. 8c**). This arrangement presents as an overall L-shape of the I3A hydrophobic cap. The hydrophobic surface of AJH-836 comprises highly exposed methyl groups of the pivaloyl R1 and the branched alkylidene R2 groups (**Ext. Fig. 8b**) that create a ridge at an ∼20° angle with the long axis of the groove. Thus, each ligand modulates the shape and the hydrophobicity of the C1Bδ membrane-binding region in its own unique way by engaging its R groups in hydrophobic interactions with the same set of protein residues (Met239, Leu250, Trp252, Leu254, V255).

The hydrophilic groups of the ligands orient towards the polar regions of the groove and recapitulate the DAG hydrogen-bonding pattern: the carbonyl oxygen (O3 in Prostratin and AJH-836; O4 in PDBu and I3A) and the hydroxyl group O1-H form hydrogen bonds with Thr242, Leu251, and Gly253 (**Fig. 4c**, red dashed lines). In addition, Prostratin and PDBu engage a second hydroxyl group (O4-H and O5-H, respectively) in hydrogen bonding with the carbonyl oxygen of Gly253. In I3A, there are two additional hydroxyls (O2-H and O5-H) that hydrogen bond to the carbonyl oxygens of Leu251 and Gly253, respectively (**Fig. 4c**, blue dashed lines). AJH-836 is the only ligand of the four with no additional ligand-protein hydrogen bonds compared to DAG.

The PDBu and Prostratin structures explain the findings of the previous structure-activity studies of phorbol ester derivatives^22,37,38^ that identified the essential role of the hydrophobic substituent, R1 at position C-12, in the PKC membrane insertion and activation. The increased potencies of 12,13-diesters with hydrophobic substituents (e.g., PDBu) relative to 12-deoxyphorbol esters (e.g., Prostratin)^37^ are due to the R1 group complementing the hydrophobic rim of loop β34 that undergoes membrane insertion. The I3A complex provides a structural rationale for the relative potencies of ingenol derivatives reported in the HIV-1 latency reversal studies^15^. The most potent ingenols exhibit conformationally restricted R1 substituents that can be accommodated in the depression formed by Gly253 and bracketed by Trp252 and Leu254. In the lactone complex, a combination of the *E* isomer and its “*sn*-1” binding mode (**Extended discussion 3**) affords a favorable arrangement of bulky R1 and R2 groups within the hydrophobic rim of C1Bδ. This configuration likely constitutes the structural basis for why AJH-836 display its marked selectivity for novel versus conventional PKC isoforms^36^.

### Comparative analyses of the C1Bδ-agonist structures

Comparative analyses of our DAG- and ligand-bound structures (**Ext. Fig. 9**) demonstrate why DAG-sensing C1 domains are capable of binding chemically diverse ligands with high affinity – a property that is driving the design of pharmacological agents. The amphiphilic binding groove, with progressively increasing hydrophobicity towards the rim of the membrane-binding region, is “tuned” to accommodate ligands with matching properties. The placement of three oxygen-containing groups, highlighted in **Ext. Fig. 9b-e**, ensures that the ligand is anchored to the polar groove regions. The ring structure invites intercalation between the C1 membrane binding loops, while the hydrophobic substituents that protrude outward from the groove, akin to the DAG acyl chains, contribute to membrane anchoring of the complex.

The comparative analysis also enables the assignment of specific functional roles to the residues of the C1 domain consensus sequence (**Ext. Fig. 9f** and **Extended discussion 1**)^29^. Two groups of residues are of particular significance. The first group consists of the four strictly conserved non-Zn^2+^ coordinating residues: Pro241, Gly253, Gln257 that directly interact with DAG (**Fig. 2b-c, 3a, Ext. Fig. 4a-b, d**); and Gly258 that ensures conformational flexibility of the β34 loop (**Extended discussion 1**). The second group comprises three hydrophobic residues: Leu250, Trp252, and Leu254 of loop β34 that show the deepest membrane insertion (**Fig. 3b**). Together with strictly conserved Pro241 and the consensus aromatic residue Phe243, these three residues form the outside hydrophobic “cage” that surrounds the various bound ligands (**Ext. Fig. 9g,h**). The spatial arrangement of the cage residues not only shields the hydrophilic ligand moiety from the hydrophobic membrane environment (**Fig. 2b,c; Fig. 4**), but also enables the loop region of C1Bδ to effectively interface with peripheral lipids. The latter function is aptly exemplified by Trp252, whose highly lipophilic indole sidechain reorients towards the loop region upon the formation of C1-ligand complexes in the membrane-mimicking environment. Given that all C1 complexes reported in this work are with potent PKC agonists, we posit that the reorientation of the Trp252 in C1Bδ (**Fig. 2a**) is an essential aspect of the mechanism of agonist capture in the membrane environment. The Trp252 conformation and interaction patterns are directly relevant to the question of DAG sensitivity of PKC isoforms – i.e. the parameter that defines the intrinsic thresholds of DAG-mediated activation (**Extended discussion 2**). Our structural data (supported by previous NMR work^23-25^) suggest that higher hydrophobicity and lipophilicity of Trp confers thermodynamic advantages and hence higher DAG-sensitivity to the C1B domains of novel PKC isoforms relative to conventional PKC isoforms that carry a Tyr at the equivalent position.

The atomistic details of our high-resolution structures of five C1Bδ-ligand complexes, particularly with regard to the arrangement of the ligand hydrophobic substituents within the binding groove and assignment of specific functional roles to the key C1 residues, provide unprecedented insight into the structural basis of DAG sensing. These also provide key information for guiding design of therapeutic agonists that selectively target proteins within the DAG effector family. Our work represents a critical advance that resolved the Gordian knot of C1-agonist crystallization. This general strategy now paves the way for the structural characterization of other C1-agonist complexes.

## Methods

### Expression, purification, and isotope enrichment of C1Bδ

The cDNA segment encoding the C1B domain of protein kinase C (PKC) δ isoenzyme from *Rattus norvegicus* (amino acids 229-281) was sub-cloned into pET SUMO expression vector (Invitrogen). The His_6_-SUMO-C1Bδ fusion protein was expressed in *Escherichia coli* BL21(DE3) Rosetta2 cells (Millipore Sigma). For the natural abundance preparations, the cells were grown in LB broth until OD_600_=0.6, followed by the induction of protein expression with 0.5 mM IPTG at 18 °C for 16 hours. For the isotopically enriched C1Bδ preparations, we used the suspension method^39^ in the M9 minimal medium supplemented with ^15^NH_4_Cl and D-^13^C_6_-glucose as nitrogen and carbon sources, respectively. To obtain [∼80% ^2^H, U-^15^N,^13^C]-enriched C1Bδ, M9 was prepared in 100% D_2_O and additionally supplemented with 1 g of ^15^N,^13^C,^2^H ISOGRO^®^ (Sigma). Cell harvesting, lysis, and C1Bδ purification were carried out as previously described^23,24^. The purified protein was stored at 4 °C in the “storage buffer” comprising 50 mM MES at pH 6.5, 150 mM KCl, and 1 mM TCEP, until further use. For NMR experiments, C1Bδ was exchanged into an “NMR buffer” composed of 20 mM d_4_-Imidazole at pH 6.5, 50 mM KCl, 0.1 mM TCEP, 0.02% NaN_3_, and 8% D_2_O.

### Preparation of isotropically tumbling bicelles

Chloroform solutions of long-chain 1,2-dimyristoyl-*sn*-glycero-3-phosphocholine (DMPC) and short-chain 1,2-dihexanoyl-*sn*-glycero-3-phosphocholine (DHPC) (Avanti Polar Lipids), or their deuterated versions d_54_-DMPC (Avanti Polar Lipids) and d_40_-DHPC (Cambridge Isotope Laboratories), were aliquoted and dried extensively under vacuum. The bicelles of q=0.5 (defined by the DMPC to DHPC molar ratio of 1:2) were prepared by suspending the dried lipid films in the NMR buffer, as previously described^40^. Additional lipid components: 1,2-dimyristoyl-*sn*-glycero-3-phospho-L-serine (DMPS) and di-octanoyl-*sn*-1,2-glycerol (DAG) were incorporated for all DAG-binding experiments, to produce the final molar ratios of DMPC:DMPS:DAG=75:15:10. For the paramagnetic relaxation enhancement (PRE) measurements, 1-palmitoyl-2-stearoyl-(14-doxyl)-*sn*-glycero-3-phosphocholine (14-doxyl PC) was incorporated into bicelles to give on average ∼1 molecule per leaflet. DMPS, DAG, and 14-doxyl PC were obtained from Avanti Polar Lipids. The lipid concentrations of final bicelle preparations were measured using the phosphate determination assay^41^.

### NMR spectroscopy

All NMR experiments were carried out at 25 °C (calibrated with d_4_-methanol) on the Avance III HD NMR spectrometer (Bruker Biospin), operating at a ^1^H Larmor frequency of 800 MHz (18.8 T) and equipped with a cryogenically cooled probe. The data were processed with NMRPipe^42^ and analyzed with NMRFAM-Sparky^43^ distribution. The backbone amide (^15^NH) and methyl (^13^CH_3_) resonance assignments of C1Bδ were obtained from our previous work^24^ and the BMRB entry 17112^44^.

### NMR detection of C1Bδ-agonist complex formation in bicelles

The ternary C1Bδ-agonist-bicelle complexes were assembled in the “NMR buffer” by combining solutions of the isotopically enriched protein, bicelles, and agonists. DAG was incorporated at the bicelle preparation stage (*vide supra*). The 30-40 mM ligand stock solutions were prepared in d_6_-DMSO from the crystalline solids (Phorbol-12,13-dibutyrate, Prostratin, and Ingenol-3-angelate, all from Sigma-Aldrich^→^; and AJH-836, custom-synthesized in Prof. Jeewoo Lee’s laboratory). The samples for the [^15^N,^1^H] HSQC ([^13^C,^1^H] HSQC) experiments contained 0.4 mM [U-^15^N,^13^C]-enriched C1Bδ and 100 mM bicelles (0.3 mM [∼80% ^2^H, U-^15^N,^13^C]-enriched C1Bδ and 80 mM deuterated bicelles). At these concentrations, the bicelle particles are approximately equimolar to protein. The protein-to-ligand molar ratios were 1:8 (DAG), 1:1.2 (PDBu/Prostratin/Ingenol 3-angelate), and 1:6 (AJH-836). The residue-specific chemical shift perturbations (CSPs, Δ) between the apo and agonist-bound C1Bδ were calculated using the following equation:

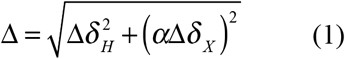

where Δ*δ*_*H*_ and Δ*δ*_*X*_ are the chemical shift changes of ^1^H and X (^15^N or ^13^C), respectively; and α=0.152 (^15^N) or 0.18 (^13^C).

### PRE and NOESY experiments

The residue-specific PRE values of ^1^H_N_ resonances, Γ_2_, were determined using the ^13^C-decoupled [^15^N,^1^H] TROSY-HSQC experiment. All data were collected in the interleaved manner, with a two-point (0 and 10 ms) relaxation delay scheme^45^. The diamagnetic ^1^H_N_ transverse relaxation rate constants, *R*_*2,dia*_, were obtained on the NMR sample comprising 0.4 mM [∼80% ^2^H, U-^15^N,^13^C]-enriched C1Bδ, 100 mM (total lipid) bicelles, and 3.2 mM DAG. The paramagnetic sample was prepared by a 1-hr room-temperature incubation of the diamagnetic sample with a dry film of 14-doxyl PC, and subsequently used to obtain the ^1^H_N_ *R*_*2,para*_ values. The error was estimated using the r.m.s.d. of the base plane noise. The Γ_2_ values were calculated using the following equation:

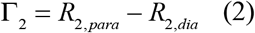

3D ^15^N-edited NOESY-TROSY experiment was carried out with the mixing time of 120 ms on a sample containing 0.4 mM [∼80% ^2^H, U-^15^N,^13^C]-enriched C1Bδ and 100 mM (total lipid) deuterated bicelles that contained 3.2 mM DAG. Inter-molecular C1Bδ-DAG ^1^H-^1^H NOEs were identified based on the available assignments of the protein amide resonances and characteristic ^1^H chemical shifts of the *sn*-1,2 stereoisomer of DAG. The DAG chemical shifts were obtained using the ^13^C-^1^H HSQC spectrum of the 0.4 mM [^13^C,C1-C3]-racemic DAG (custom synthesized by Avanti Polar Lipids) in 10 mM d_38_-dodecylphosphocholine (DPC, Sigma) micelles, and matched the literature values^46^.

### Crystallization of apo C1Bδ and its complexes with agonists

Due to a patent application pending with the United States Patent and Trademark Office, the authors have redacted this section of the methods.

### Data collection, processing, and model building of C1Bδ-agonist complexes

For the apo structure and all the ligands except AJH-836 the data were collected on home source Cu k-alpha X-ray generator. The data were indexed, scaled and integrated by PROTEUM software^47^. For the C1Bδ-DAG complex crystal, the data were collected at Argonne National Lab APS synchrotron, beamline 23ID, and for the C1Bδ-AJH-836 complex crystal – at ALS synchrotron at Berkley, beamline BL502. The data were indexed, integrated and scaled by the beamline auto-processing pipeline (XDS^48^, POINTLESS^49^, and AIMLESS^50^ software packages). Structures were solved by molecular replacement, using the PDB entry 1PTQ as a search model^19^. This was followed by iterative cycles of refinement with PHENIX.REFINE and manual building in COOT^51,52^. Polder omit maps were generated using PHENIX^53^. Ligands were created using ELBOW.BUILDER and JLIGAND^54,55^. Structural analyses were carried out using UCSF Chimera^56^, CCG Molecular Operating Environment (MOE)^57^, and LigPlot^+58^.

Although the diffraction data of the C1Bδ-DAG complex could be indexed and scaled in the F23 space group, the structure was solved in the lower symmetry H3 space group because of the non-uniform lipid molecules in the solvent channels. We have built only the lipids for which well-ordered electron density of head groups was present. The peripheral lipids and detergents are less ordered, with average B factors of 71 and 52 for DAG and DPC, respectively (average protein B factor is 36). However, there are likely more lipids and/or detergents in the solvent channels, as evidenced by the multitude of positive difference electron density peaks that are larger than water, some reaching across the symmetry axis.

The coordinates of all structures were deposited in the Protein Data Bank. The accession numbers and statistics are given in the **Supporting information Tables 1 and 2**.

### Data availability

The atomic coordinates and structure factors are deposited in the PDB (https://www.rcsb.org) under the accession codes (Ligand) as: 7KND (apo), 7L92 (DAG), 7LEO (AJH-836), 7LF3 (AJH-836), 7KNJ (PDBu), 7LCB (Prostratin), and 7KO6 (I3A). Relevant data supporting the findings of this work are available within the manuscript. Requests for further information should be addressed to the corresponding author.

## Acknowledgements

This research was supported by the U.S. National Institute of Health grant R01 GM108998 and Texas A&M institutional funds to T.I.I., and Welch Foundation grant A-0015 to J.C.S. We are grateful to Dr. Vytas A. Bankaitis for critical reading of the manuscript and suggestions.

## Author contributions

S.S.K. and T.I.I. conceptualized the work, developed the crystallization approach, and designed NMR experiments. S.S.K. prepared all samples, crystallized the complexes, and collected and analyzed NMR data. I.V.K. and S.S.K. collected the X-ray diffraction data. I.V.K. built and refined the models. S.S.K., I.V.K., J.C.S., and T.I.I. analyzed the structural data. J.A. and J.L. synthesized and provided the AJH-836 compound. T.I.I. and S.S.K. wrote the manuscript with input from all coauthors. T.I.I. supervised the project.

## Supplementary information

This manuscript is accompanied by supplementary information containing referenced extended discussion and crystallography statistics tables.

## Extended Data: Figures

**Extended data Fig.1.**
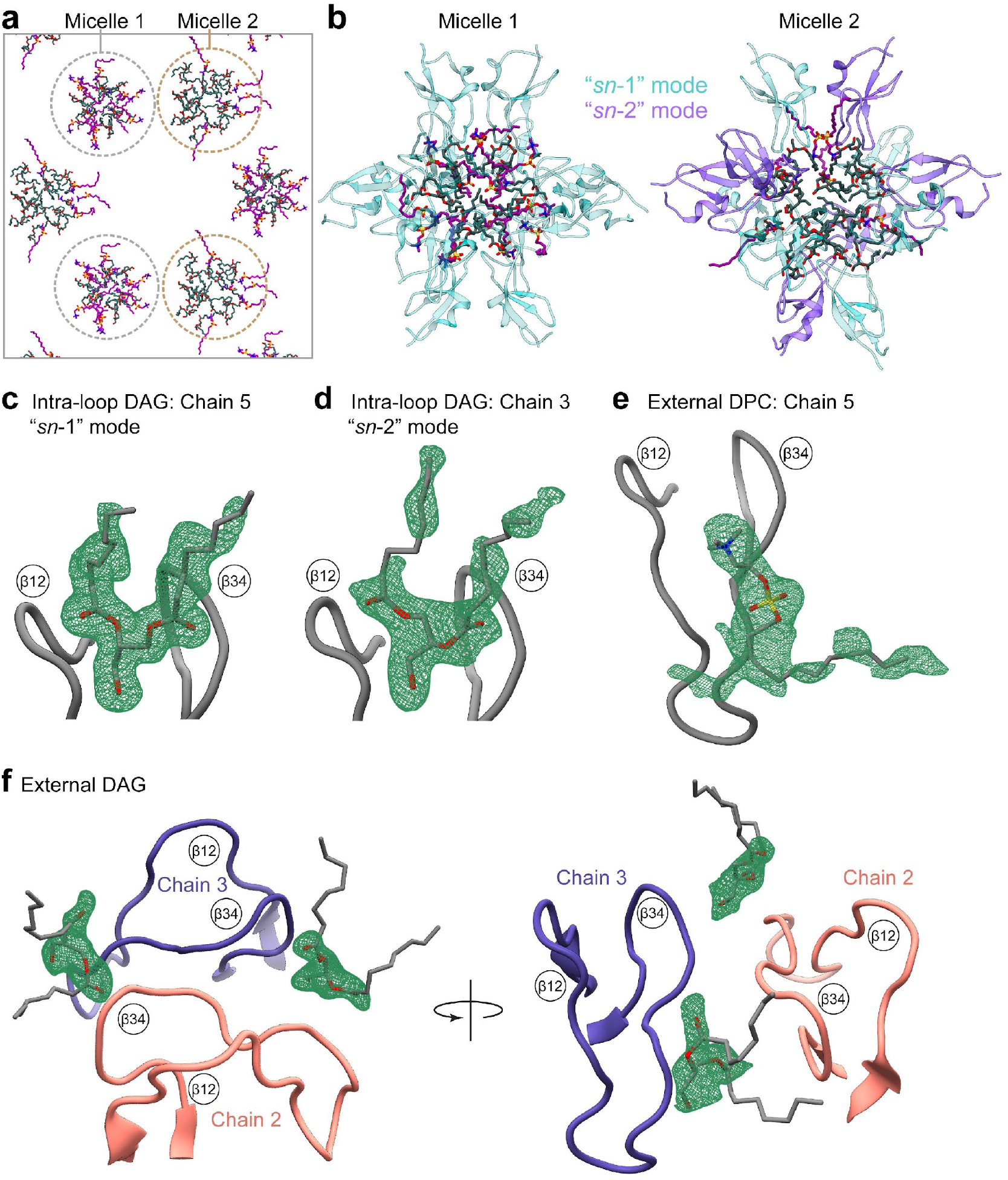
Organization of DAG and DPC molecules in the crystal. **a**, The DAG and DPC molecules are ordered within the crystal lattice as two mixed micelles, micelle 1 (12 DAG and 12 DPC molecules) and micelle 2 (18 DAG and 6 DPC molecules). **b**, The C1Bδ chains are arranged with their lipid-sensing loops directly binding to DAG within micelles. Micelle 1 supports only “*sn-1*” binding mode, while micelle 2 supports both, “*sn-1*” and “*sn-2*”. **c-f**, Representative 2*F*_*0*_-*F*_*c*_ Polder omit electron density maps of DAG and DPC molecules. **c**, and **d**, The 2*F*_*0*_-*F*_*c*_ electron density maps of DAG molecules bound in the “*sn*-1” (chain 5, contoured at 2.7σ) and “*sn*-2” (chain 3, contoured at 2.8σ) modes within the intra-loop groove of C1Bδ. **e**, The 2*F*_*0*_-*F*_*c*_ electron density map, contoured at 2.8σ, of the DPC bound externally to chain 5 of C1Bδ. **f**, The 2*F*_*0*_-*F*_*c*_ electron density maps, each contoured at 2.5σ, of two DAG molecules bound externally to C1Bδ chains 2 and 3.

**Extended data Fig.2.**
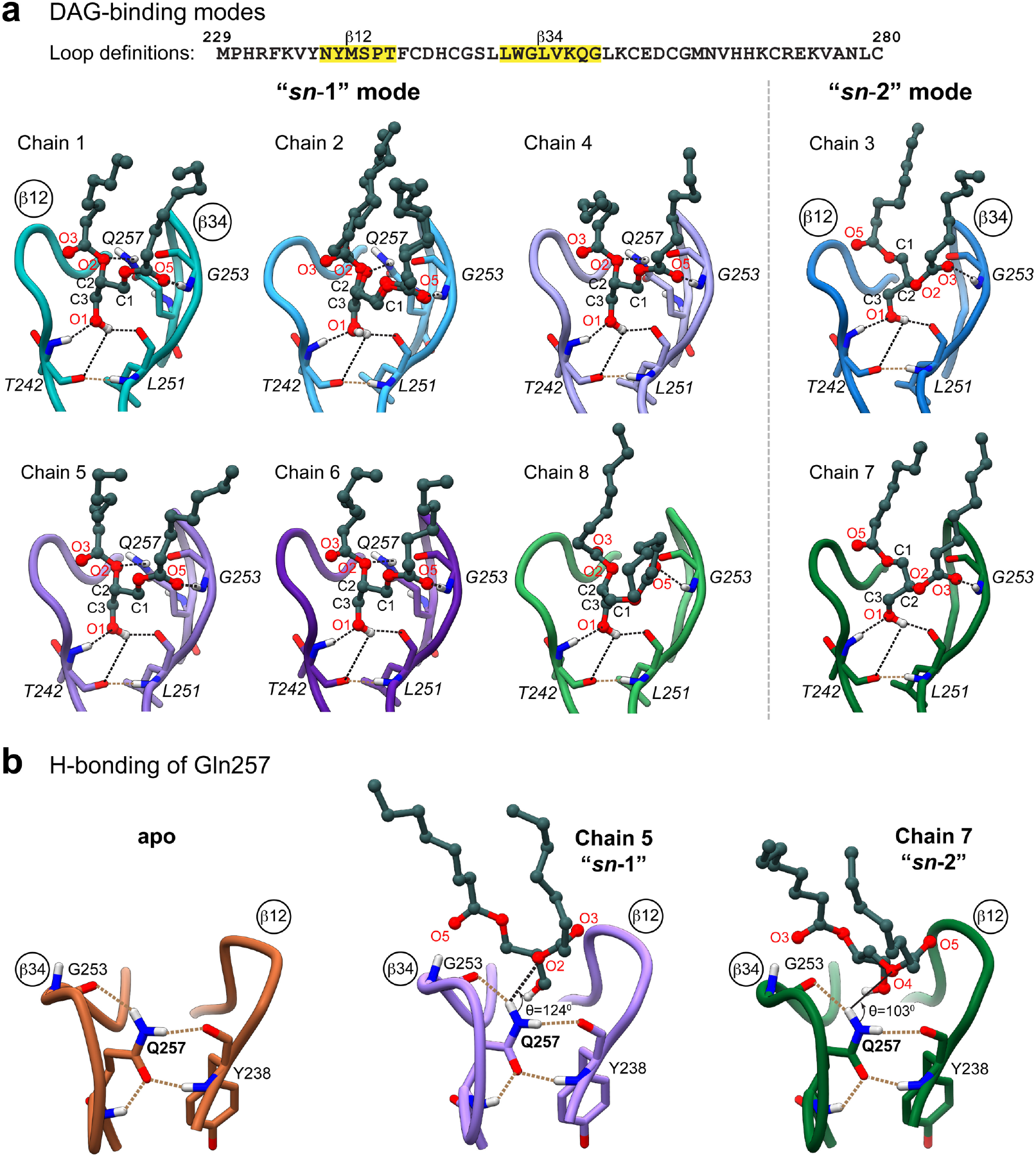
DAG binding modes and hydrogen-bonding patterns for the C1Bδ protein chains in the asymmetric unit. **a**, 6 and 2 C1Bδ protein chains have DAG bound in the “*sn*-1” and “*sn*-2” modes, respectively. In the “*sn*-1” mode, the H-bond acceptor of Gly253 amide hydrogen is the carbonyl oxygen of the *sn*-1 ester group (O5), whereas in the “*sn*-2” mode the acceptor is the carbonyl oxygen of the *sn*-2 ester group (O3). The sidechain of Gln257 makes a hydrogen bond with the alkoxy O2 oxygen of DAG in 5 out of 6 “*sn*-1” complexes. **b**, Gln257 stabilizes the intra-loop region by forming four hydrogen bonds (brown dashed lines) that stitch the β12 and β34 loops together. These bonds are present in the apo and all DAG-complexed C1Bδ structures. The hydrogens of the Gln257 sidechain amide group also serve as the H-bond donors to the DAG alkoxy oxygen O2 in the “*sn*-1” mode (illustrated using Chain 5, black dashed line). The equivalent interaction in the “*sn*-2” mode involving the alkoxy O4 oxygen would have a non-optimal angle and therefore is unlikely to occur.

**Extended data Fig.3.**
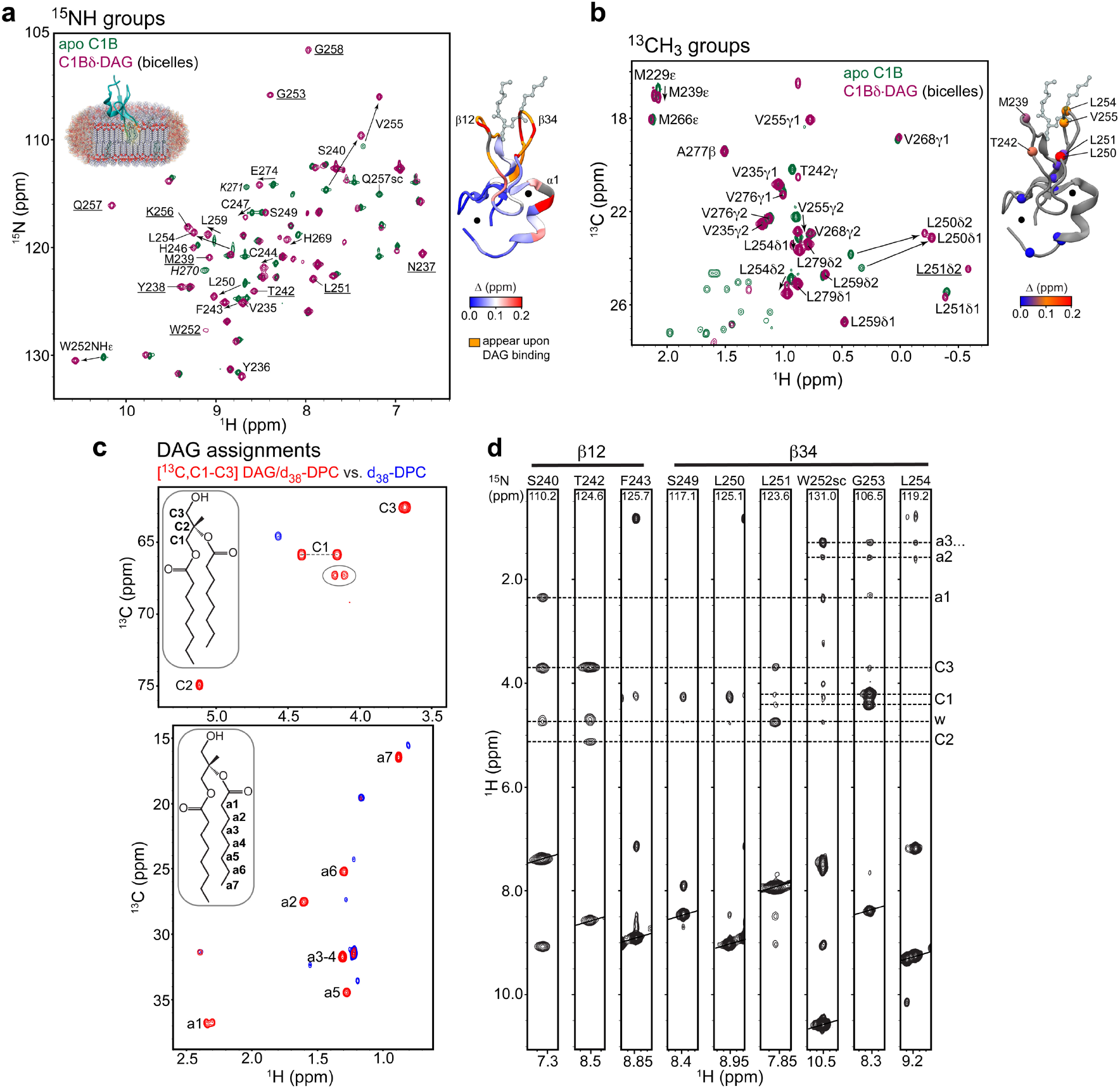
Formation of the ternary C1Bδ-DAG-bicelle complex detected by solution NMR spectroscopy. Overlays of the ^15^N-^1^H (**a**) and the methyl ^13^C-^1^H HSQC spectra (**b**) of apo C1Bδ (green) and DAG-bound complex in bicelles (maroon). The formation of the C1Bδ-DAG-bicelle complex is evidenced by: (i) the chemical shift changes of the amide and methyl peaks mapped onto the 3D structure of C1Bδ (right side of panels **a** and **b**), and (ii) the reappearance, upon DAG binding, of the amide cross-peaks (underlined, and mapped on 3D structure in orange) that were exchange-broadened in apo C1Bδ. In (**a**), two residues that belong to the C-terminal α-helix, K271 and H270 (italicized), broaden upon association with bicelles and their cross-peaks are not visible at this contour threshold. **c**, Atom-specific assignments of the ^13^C and ^1^H resonances of DAG mapped onto the ^13^C-^1^H HSQC spectra of DAG embedded into the deuterated DPC micelles (red). Circled cross-peaks belong to the *sn*-1,3 isomer. The spectrum of residual ^1^H in deuterated DPC (blue) is shown as an overlay for comparison. **d**, ^1^H-^1^H_N_ strips from the 3D ^15^N-edited NOESY-TROSY spectrum obtained for the C1Bδ-DAG-bicelle complex. The NH groups of loop residues show inter-molecular NOEs to the ^1^H atoms of the DAG glycerol moiety that are consistent with the “*sn*-1” binding mode. Also present are the NOEs to the methylene ^1^H that belong to the acyl groups of DAG or bicelle lipids. “w” denotes water protons. Intra-protein ^1^H-^1^H NOEs are suppressed by protein deuteration.

**Extended data Fig.4.**
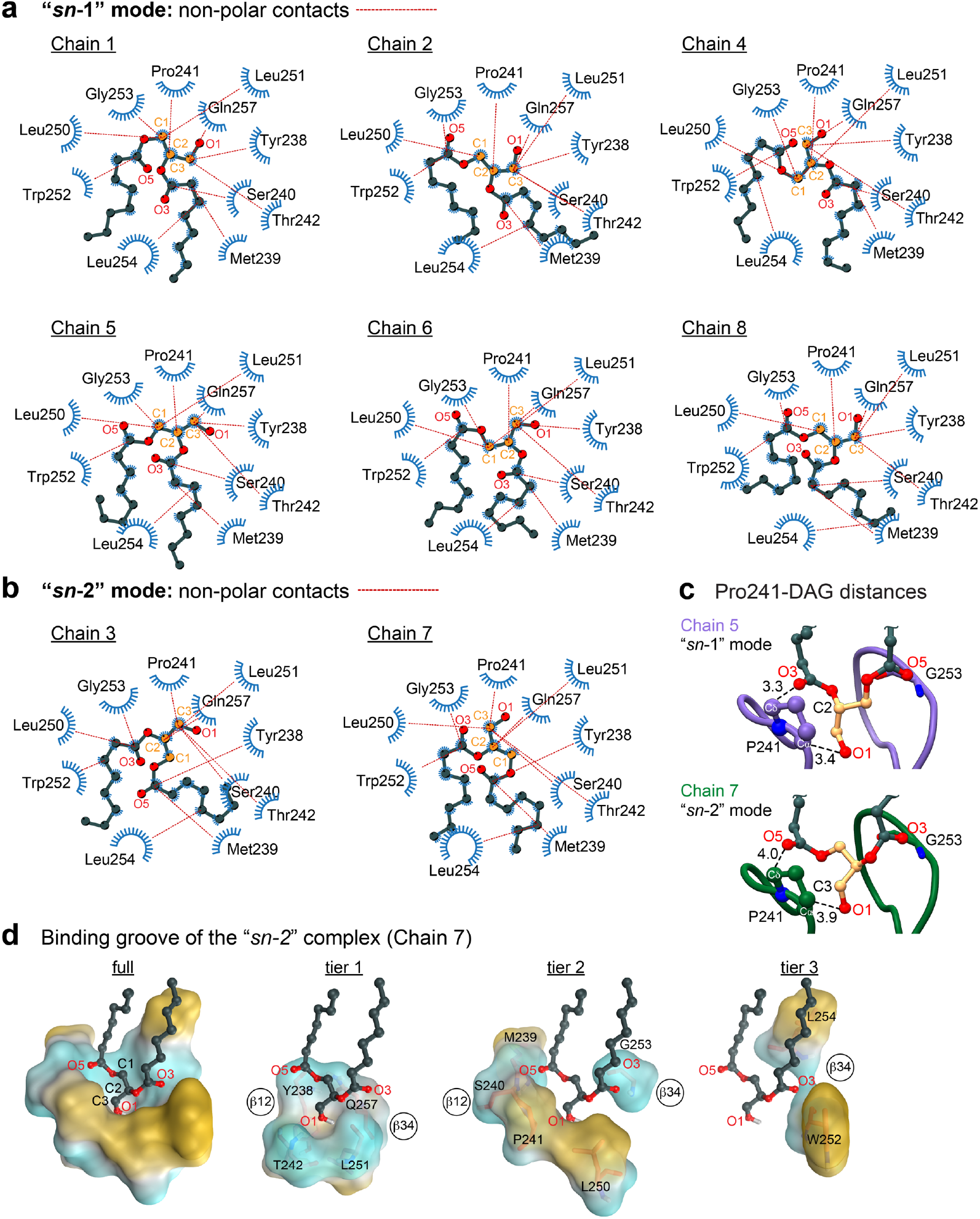
Nonpolar C1Bδ-DAG contacts and the properties of the DAG-binding groove in the “*sn*-2” complex. **a and b**, 2D LigPlot^+^ diagrams of nonpolar C1Bδ-DAG interactions. The contact cutoff for backbone and sidechain carbon atoms is 4.5 Å. A subset of contacts is shown with lines to guide the eye. **c**, Pro241 makes non-polar contacts with C2 (“*sn-*1” mode) or C3 (“*sn-*2” mode) of DAG. In addition, pyrrolidine Cδ/Cα are within optimal distance from DAG oxygens in both modes to support C-H…O interactions (denoted by dashed lines with distances) **d**, Polar backbone atoms and hydrophobic sidechains of DAG-interacting C1Bδ residues create a binding site whose properties are tailored to capture the amphiphilic DAG molecule. This is illustrated through the deconstruction of the binding site into three tiers that accommodate the glycerol backbone (tier 1), the *sn*-1/2 ester groups (tier 2), and the acyl chain methylenes (tier 3).

**Extended data Fig.5.**
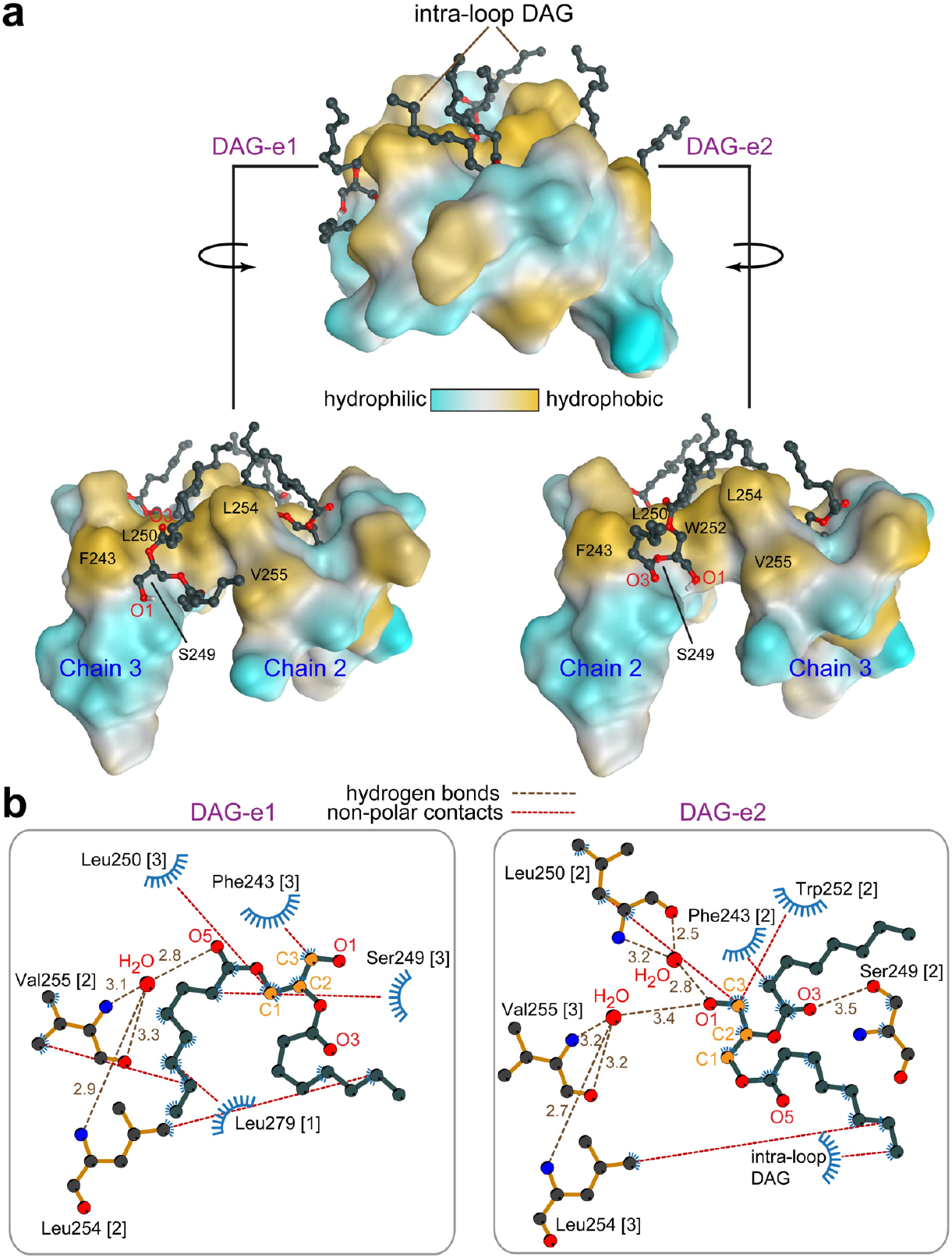
Peripheral interactions of DAG with C1Bδ. **a**, Two DAG molecules, DAG-e1 and DAG-e2, peripherally bind to the external hydrophobic grooves created by the C1Bδ chains 2 and 3. **b**, The peripheral C1Bδ-DAG interactions involve hydrogen bonds and non-polar contacts, as shown in the 2D LigPlot^+^ interaction diagrams. Most hydrogen bonds are mediated by water molecules (red spheres) that bridge the polar groups of external DAG molecules to the polar backbone atoms of Val255, Leu254, and Leu250. Non-polar contacts involve hydrophobic residues Phe243, Leu250, Trp252, Leu254, and Val255. The protein chain number is given in square brackets.

**Extended data Fig.6.**
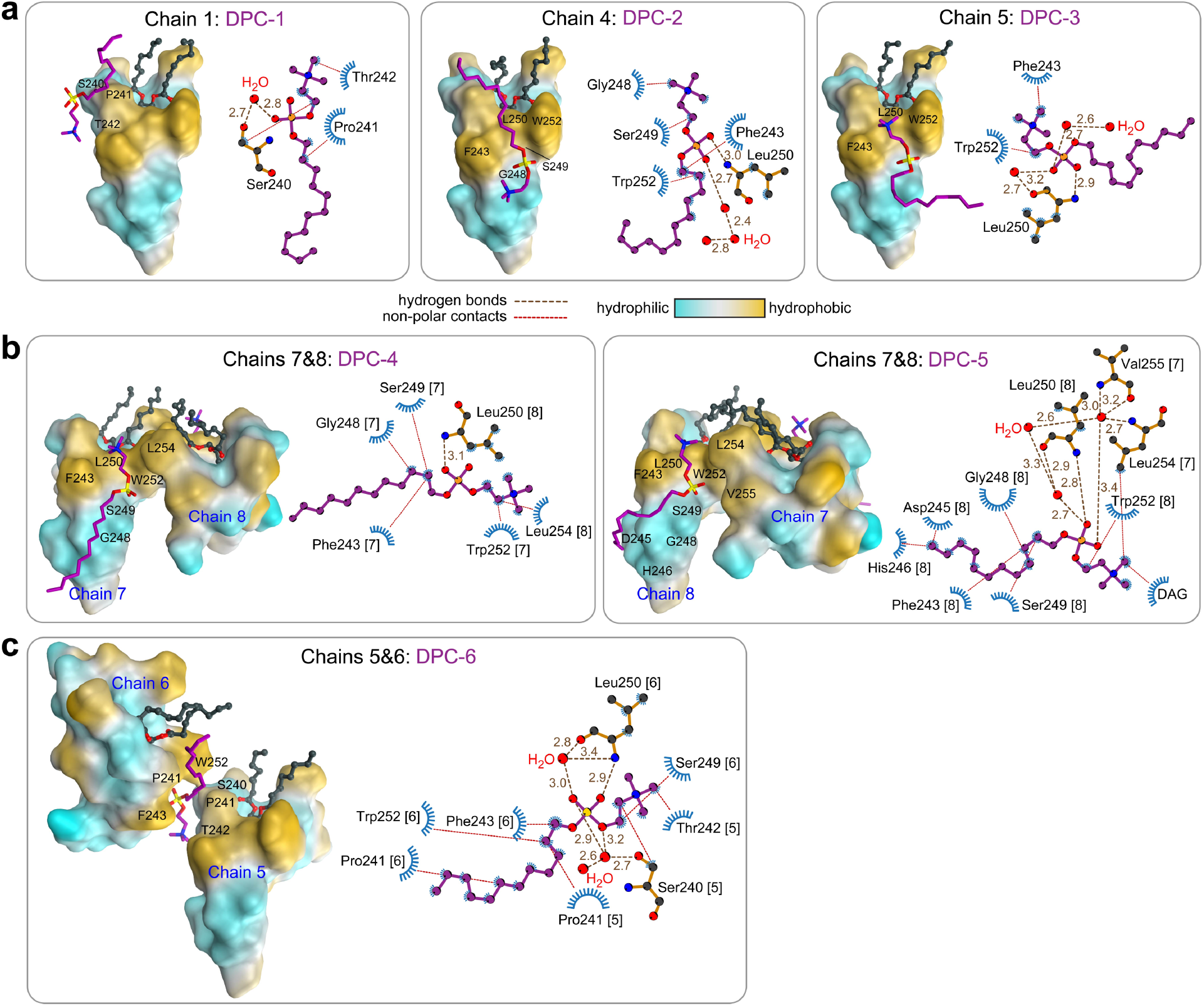
Peripheral interactions of DPC with C1Bδ. The 8-chain asymmetric unit contains six DPC molecules that peripherally interact with C1Bδ. **a**, DPC-1, -2, and -3 associate with individual C1Bδ chains 1, 4, and 5. **b**, DPC-4 and DPC-5 bind in the grooves created by chains 7 and 8. **c**, DPC-6 is wedged between chains 5 and 6. 2D interaction diagrams between C1Bδ and DPC reveal several common patterns. The phosphate oxygens of DPCs engage in hydrogen bonds with solvent molecules and the NH group of the Leu250 backbone. The methyls of the phosphocholine group, along with the aliphatic tails of DPC, form non-polar contacts with the hydrophobic residues of the β12 and β34 loops. The loop residues that show most interactions with the DPC molecules are Phe243 of β12, and Leu250 and Trp252 of β34. In (**a**)-(**c**), DPC is shown in purple, DAG in gray, and the surface of the protein chains is color-coded according to its hydrophilicity/hydrophobicity.

**Extended data Fig.7.**
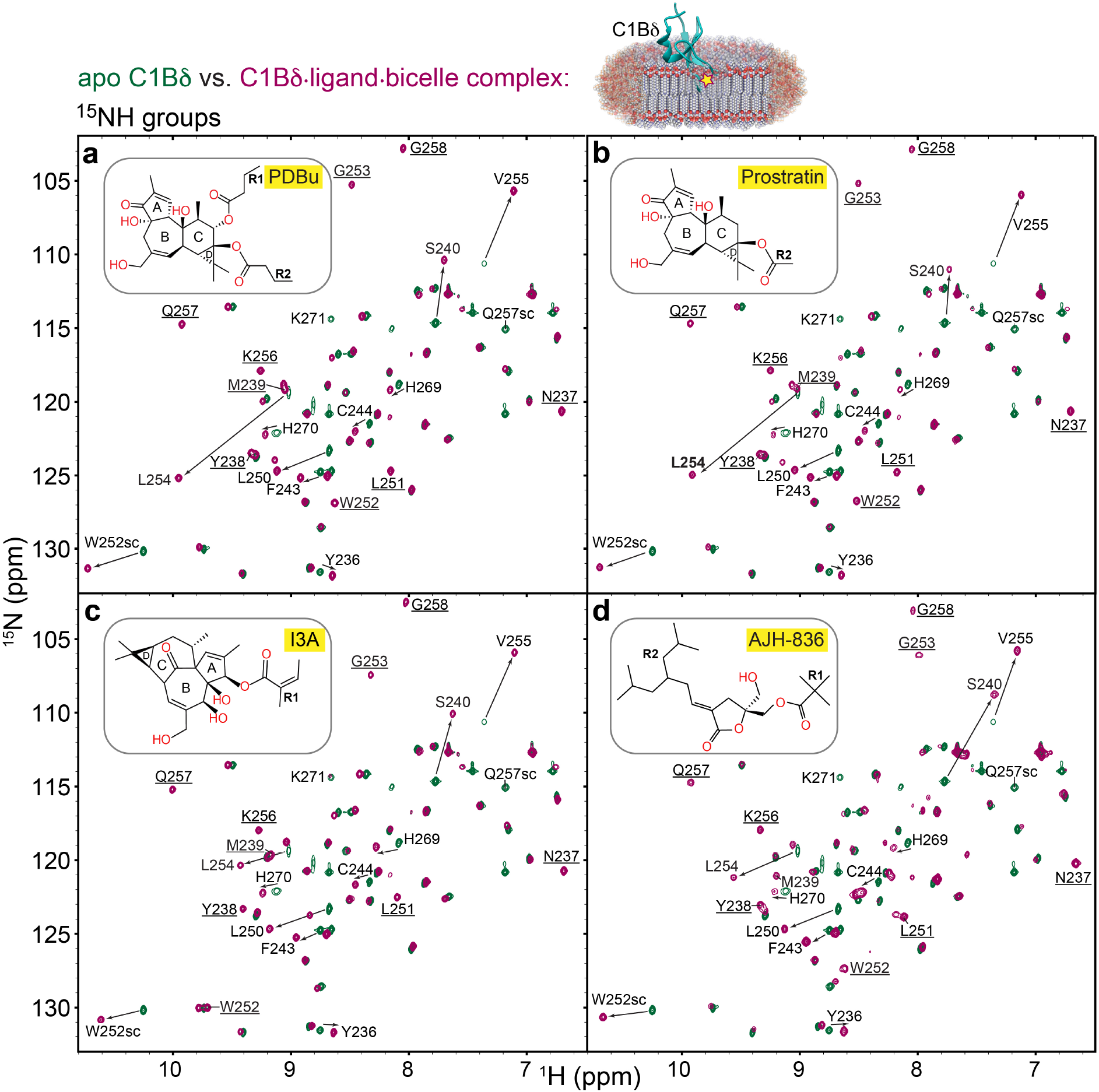
Formation of the ternary C1Bδ-ligand-bicelle complexes detected by solution NMR spectroscopy. Overlays of the ^15^N-^1^H HSQC spectra of apo C1Bδ (green) and its complexes (maroon) with **a**, PDBu; **b**, Prostratin; **c**, I3A; and **d**, AJH-836, all in bicelles. The formation of the C1Bδ-ligand-bicelle complexes is evidenced by: (i) the chemical shift changes of the amide peaks, and (ii) the appearance, upon ligand binding, of cross-peaks (underlined) that were exchanged-broadened in apo C1Bδ. The protein-to-ligand ratios were 1:1.2 (PDBu), 1:1.2 (Prostratin), 1:1.2 (I3A), and 1:6 (AJH-836).

**Extended data Fig.8.**
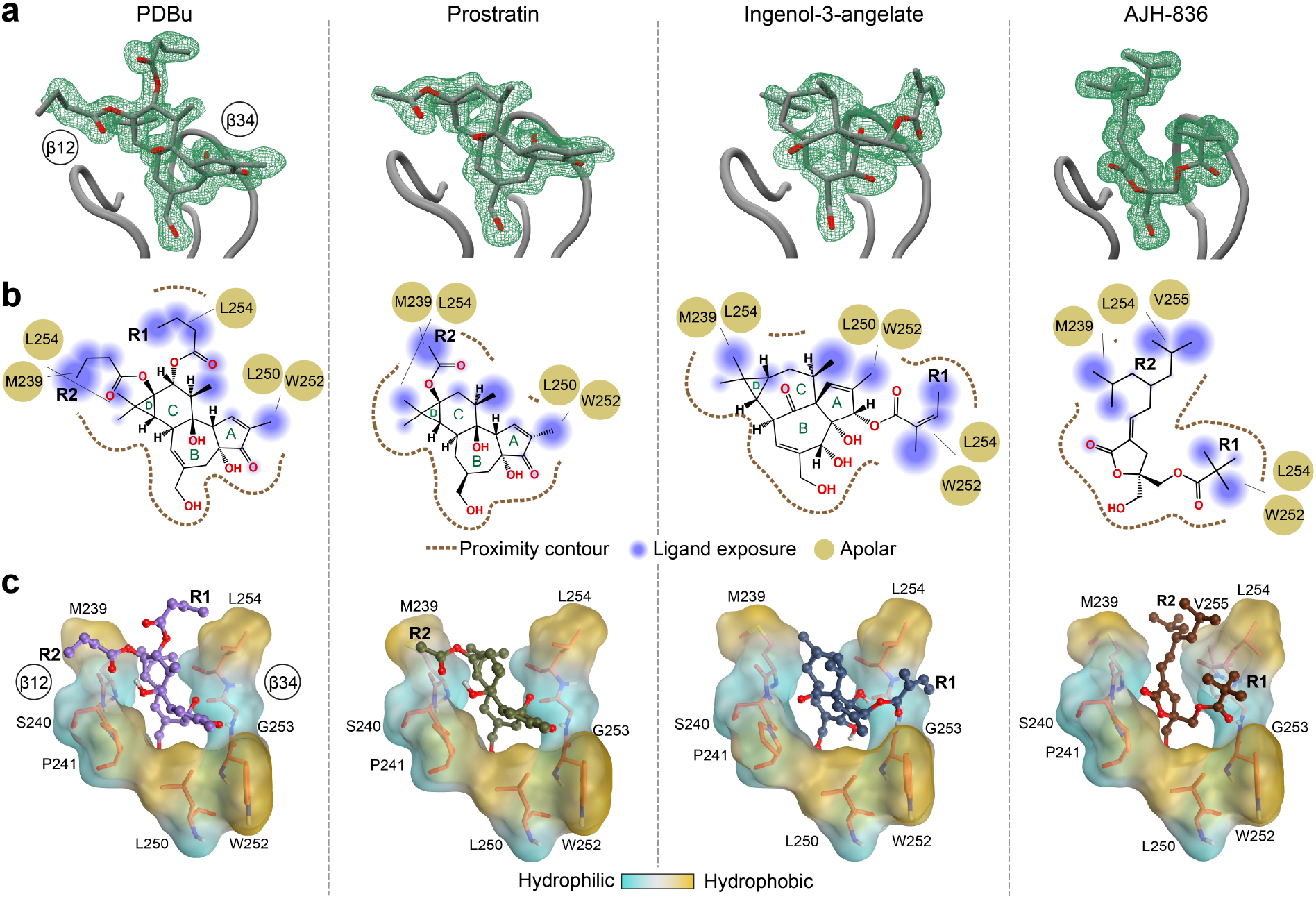
Ligand position and interactions in the groove region of C1Bδ. **a**, The 2*F*_*0*_-*F*_*c*_ Polder omit electron density maps showing well-resolved electron densities of the ligands in the C1Bδ complexes. The maps were contoured at 2.0σ (PDBu), 2.8σ (prostratin), and 2.3σ (ingenol-3-angelate and AJH-836). **b**, 2D diagrams of the ligand placement in the binding groove outlined with the dashed line. The ligand chemical groups that protrude out of the groove are highlighted with blue spheres, with the radius proportional to the degree of solvent exposure. Also shown are protein residues involved in non-polar interactions with the hydrophobic moieties of ligands. **c**, 3D representations of the ligand-filled grooves that show the placement of hydrophobic R groups in the complexes.

**Extended data Fig.9.**
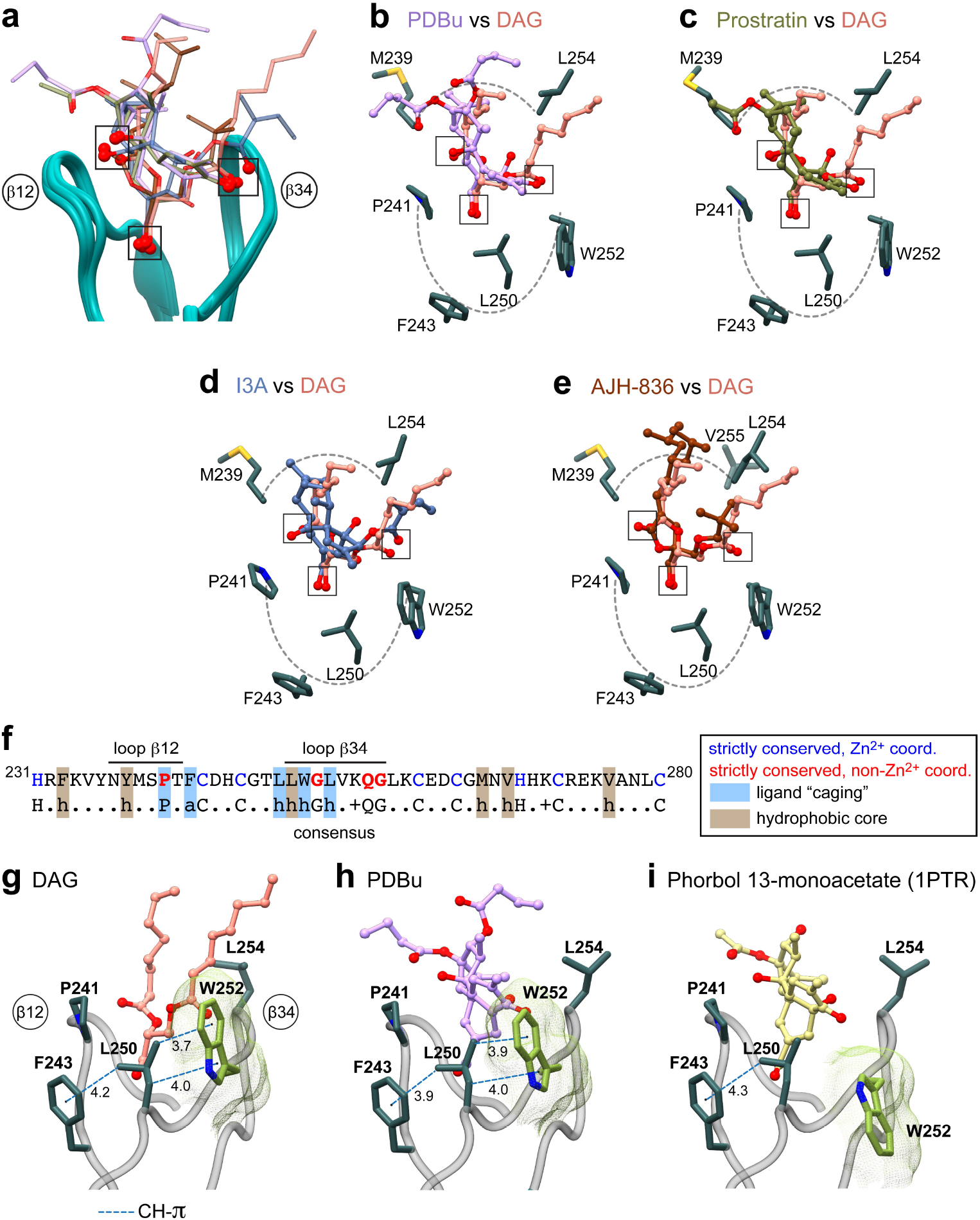
Structural analysis of C1Bδ-agonist complexes identifies three key oxygen-containing groups and the roles of conserved hydrophobic residues. **a**, Loop region of the backbone-superimposed C1Bδ complexes (pairwise r.m.s.d. <0.6 Å relative to chain 5 of the “*sn*-1” DAG complex). Oxygen-containing functional groups involved in the interactions with C1Bδ are highlighted by squares. **b-e**, Pairwise comparison of the binding poses of **b**, PDBu; **c**, prostratin; **d**, ingenol 3-angelate; and **e**, AJH-836 relative to that of DAG in the binding groove. Hydrophobic sidechains that envelope the ligands and form the rim of the membrane-binding region are also shown. **f**, Amino acid sequence of C1Bδ and the consensus sequence of DAG-sensitive C1 domains. **g-i**, A subset of conserved hydrophobic residues that form a “cage”-like arrangement around the ligands, with a potential to form CH-π interactions in addition to the apolar contacts. The rotameric flip of Trp252, illustrated using DAG (**g**) and PDBu (**h**) complexes, is essential for creating a contiguous hydrophobic surface. **i**, The Trp252 sidechain remains in its apo-state rotameric conformation in the complex of C1Bδ with a non PKC-activating phorbol ester that was crystallized in the absence of lipids/detergents.

**Extended data Fig.10.**
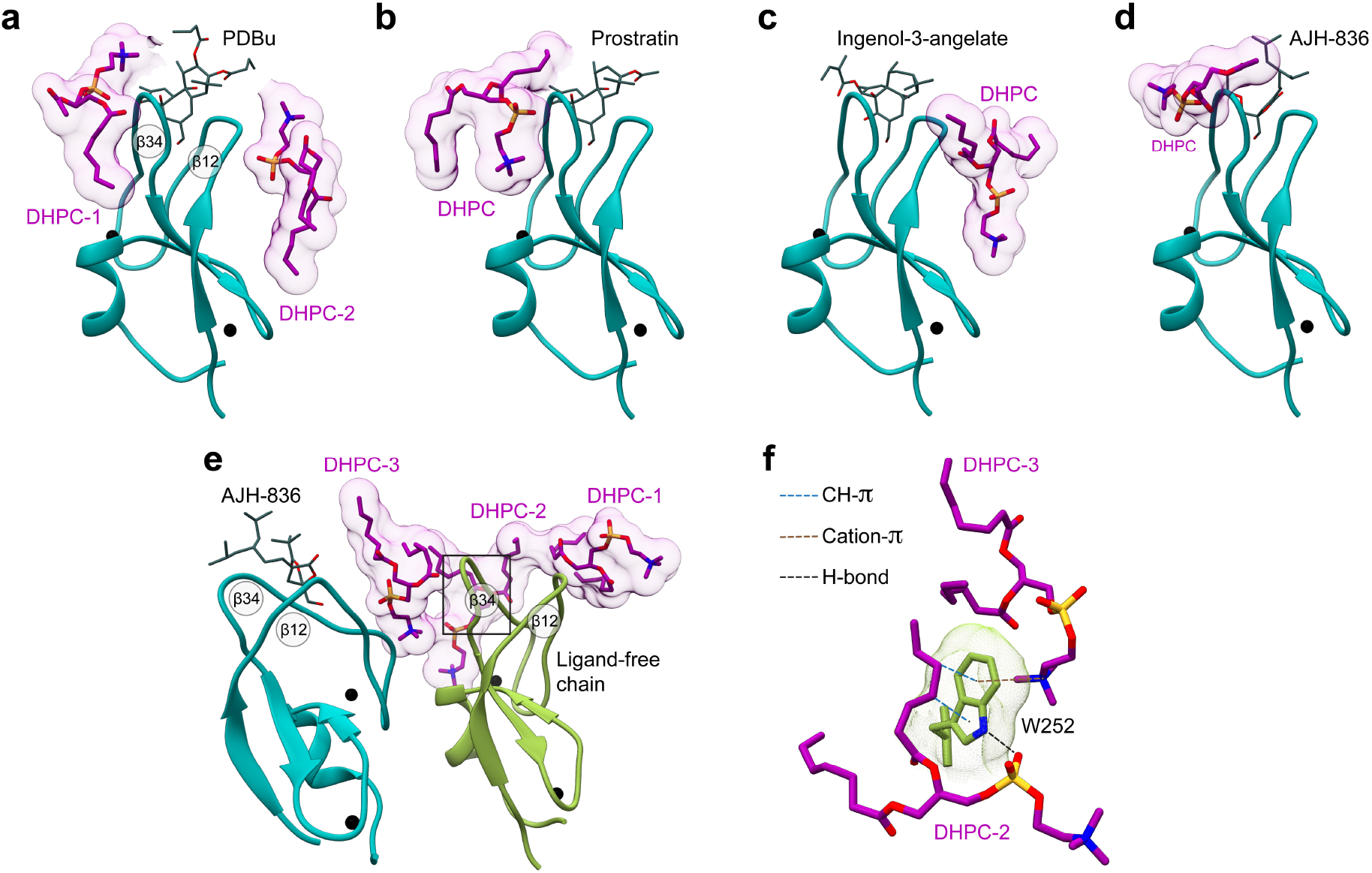
Peripheral DHPC molecules in the C1Bδ-ligand complexes. DHPC molecules peripherally associate with the membrane-binding loop regions of C1Bδ complexed to **a**, PDBu; **b**, prostratin; **c**, ingenol-3-angelate, **d**, AJH-836 (one molecule per AU), and **e**, AJH-836 (two molecules per AU). In **e**, one protein chain is ligand-free (color-coded green) and has three DHPC molecules, labeled 1 through 3 that cap the membrane-binding loop region. **f**, The versatility of potential Trp252-lipid interactions, exemplified by the Trp252 sidechain from the ligand-free C1Bδ monomer (**e**, green). In addition to non-polar contacts with the hydrophobic lipid moieties, the Trp sidechains can engage in H-bonding, cation-π, and CH-π interactions.

## References

1 Eichmann, T. O. & Lass, A. DAG tales: the multiple faces of diacylglycerol—stereochemistry, metabolism, and signaling. Cell. Mol. Life Sci. 72, 3931–3952, (2015).

2 Carrasco, S. & Mérida, I. Diacylglycerol, when simplicity becomes complex. Trends Biochem. Sci. 32, 27–36, (2007).

3 Takai, Y., Kishimoto, A., Inoue, M. & Nishizuka, Y. Studies on a cyclic nucleotide-independent protein kinase and its proenzyme in mammalian tissues. I. Purification and characterization of an active enzyme from bovine cerebellum. J. Biol. Chem. 252, 7603–7609, (1977).

4 Igumenova, T. I. Dynamics and membrane interactions of protein kinase C. Biochemistry 54, 4953–4968, (2015).

5 Protein Kinase C in Cancer Signaling and Therapy. Anticancer Res. 30, 5263, (2010).

6 Ma, Q., Gabelli, S. B. & Raben, D. M. Diacylglycerol kinases: Relationship to other lipid kinases. Advances in biological regulation 71, 104–110, (2019).

7 Gutierrez-Uzquiza, A. et al. Coordinated activation of the Rac-GAP beta2-chimaerin by an atypical proline-rich domain and diacylglycerol. Nat Commun 4, 1849, (2013).

8 Ebinu, J. O. et al. RasGRP, a Ras guanyl nucleotide-releasing protein with calcium-and diacylglycerol-binding motifs. Science 280, 1082–1086, (1998).

9 Zhao, Z. & Manser, E. Myotonic dystrophy kinase-related Cdc42-binding kinases (MRCK), the ROCK-like effectors of Cdc42 and Rac1. Small GTPases 6, 81–88, (2015).

10 Xu, J. et al. Mechanistic insights into neurotransmitter release and presynaptic plasticity from the crystal structure of Munc13-1 C1C2BMUN. eLife 6, e22567, (2017).

11 Rosse, C. et al. PKC and the control of localized signal dynamics. Nature Reviews Molecular Cell Biology 11, 103–112, (2010).

12 Castagna, M. et al. Direct Activation of Calcium-Activated, Phospholipid-Dependent Protein-Kinase by Tumor-Promoting Phorbol Esters. J. Biol. Chem. 257, 7847–7851, (1982).

13 Ly, C. et al. Bryostatin 1 Promotes Synaptogenesis and Reduces Dendritic Spine Density in Cortical Cultures through a PKC-Dependent Mechanism. ACS Chemical Neuroscience 11, 1545–1554, (2020).

14 Sloane, J. L. et al. Prodrugs of PKC modulators show enhanced HIV latency reversal and an expanded therapeutic window. Proceedings of the National Academy of Sciences 117, 10688–10698, (2020).

15 Spivak, A. M. et al. Synthetic Ingenols Maximize Protein Kinase C-Induced HIV-1 Latency Reversal. Antimicrob. Agents Chemother. 62, (2018).

16 Hardman, C. et al. Synthesis and evaluation of designed PKC modulators for enhanced cancer immunotherapy. Nat Commun 11, 1879, (2020).

17 Nakagawa, Y. et al. A Simple Analogue of Tumor-Promoting Aplysiatoxin Is an Antineoplastic Agent Rather Than a Tumor Promoter: Development of a Synthetically Accessible Protein Kinase C Activator with Bryostatin-like Activity. J. Am. Chem. Soc. 131, 7573–7579, (2009).

18 Katti, S. & Igumenova, T. I. Structural insights into C1-ligand interactions: Filling the gaps by in silico methods. Advances in Biological Regulation 79, 100784, (2021).

19 Zhang, G. G., Kazanietz, M. G., Blumberg, P. M. & Hurley, J. H. Crystal-Structure of the Cys2 Activator-Binding Domain of Protein-Kinase C-Delta in Complex with Phorbol Ester. Cell 81, 917–924, (1995).

20 Zhang, G. & Hurley, J. H. Crystallization of the protein kinase Cdelta C1B domain. Methods Mol Biol 233, 299–304, (2003).

21 Rahman, G. M. et al. Identification of the activator-binding residues in the second cysteine-rich regulatory domain of protein kinase Ctheta (PKCtheta). Biochem. J. 451, 33–44, (2013).

22 Ryckbosch, S. M., Wender, P. A. & Pande, V. S. Molecular dynamics simulations reveal ligand-controlled positioning of a peripheral protein complex in membranes. Nature Communications 8, 6, (2017).

23 Stewart, M. D., Morgan, B., Massi, F. & Igumenova, T. I. Probing the determinants of diacylglycerol binding affinity in the C1B domain of protein kinase Calpha. J. Mol. Biol. 408, 949–970, (2011).

24 Stewart, M. D., Cole, T. R. & Igumenova, T. I. Interfacial partitioning of a loop hinge residue contributes to diacylglycerol affinity of conserved region 1 domains. J. Biol. Chem. 289, 27653–27664, (2014).

25 Stewart, M. D. & Igumenova, T. I. Toggling of Diacylglycerol Affinity Correlates with Conformational Plasticity in C1 Domains. Biochemistry 56, 2637–2640, (2017).

26 Dries, D. R., Gallegos, L. L. & Newton, A. C. A single residue in the C1 domain sensitizes novel protein kinase C isoforms to cellular diacylglycerol production. J. Biol. Chem. 282, 826–830, (2007).

27 Antal, C. E., Violin, J. D., Kunkel, M. T., Skovso, S. & Newton, A. C. Intramolecular conformational changes optimize protein kinase C signaling. Chem. Biol. 21, 459–469, (2014).

28 Sigano, D. M. et al. Differential binding modes of diacylglycerol (DAG) and DAG lactones to protein kinase C (PK-C). J. Med. Chem. 46, 1571–1579, (2003).

29 Hurley, J. H., Newton, A. C., Parker, P. J., Blumberg, P. M. & Nishizuka, Y. Taxonomy and function of C1 protein kinase C homology domains. Protein Sci. 6, 477–480, (1997).

30 Kazanietz, M. G. et al. Residues in the second cysteine-rich region of protein kinase C delta relevant to phorbol ester binding as revealed by site-directed mutagenesis. J. Biol. Chem. 270, 21852–21859, (1995).

31 Choi, Y. et al. Conformationally Constrained Analogues of Diacylglycerol (DAG). 28. DAG-dioxolanones Reveal a New Additional Interaction Site in the C1b Domain of PKCδ. J. Med. Chem. 50, 3465–3481, (2007).

32 Abel, E. L., Angel, J. M., Kiguchi, K. & DiGiovanni, J. Multi-stage chemical carcinogenesis in mouse skin: fundamentals and applications. Nat Protoc 4, 1350–1362, (2009).

33 Melander, C. & Margolis, D. M. Forcing an enemy into the open. Nature Chemistry 4, 692–693, (2012).

34 Wender, P. A., Kee, J.-M. & Warrington, J. M. Practical Synthesis of Prostratin, DPP, and Their Analogs, Adjuvant Leads Against Latent HIV. Science 320, 649, (2008).

35 Hanke, C. W. et al. Efficacy and safety of ingenol mebutate gel in field treatment of actinic keratosis on full face, balding scalp, or approximately 250 cm(2) on the chest: A phase 3 randomized controlled trial. J Am Acad Dermatol 82, 642–650, (2020).

36 Cooke, M. et al. Characterization of AJH-836, a diacylglycerol-lactone with selectivity for novel PKC isozymes. J. Biol. Chem. 293, 8330–8341, (2018).

37 Kazanietz, M. G., Krausz, K. W. & Blumberg, P. M. Differential irreversible insertion of protein kinase C into phospholipid vesicles by phorbol esters and related activators. J. Biol. Chem. 267, 20878–20886, (1992).

38 Bertolini, T. M., Giorgione, J., Harvey, D. F. & Newton, A. C. Protein kinase C translocation by modified phorbol esters with functionalized lipophilic regions. J Org Chem 68, 5028–5036, (2003).

## Methods References

39 Marley, J., Lu, M. & Bracken, C. A method for efficient isotopic labeling of recombinant proteins. J. Biomol. NMR 20, 71–75, (2001).

40 Katti, S., Nyenhuis, S. B., Her, B., Cafiso, D. S. & Igumenova, T. I. Partial Metal Ion Saturation of C2 Domains Primes Synaptotagmin 1-Membrane Interactions. Biophys. J. 118, 1409–1423, (2020).

41 King, E. J. The colorimetric determination of phosphorus. Biochem. J. 26, 292–297, (1932).

43 Delaglio, F. et al. NMRPipe: a multidimensional spectral processing system based on UNIX pipes. J. Biomol. NMR 6, 277–293, (1995).

43 Lee, W., Tonelli, M. & Markley, J. L. NMRFAM-SPARKY: enhanced software for biomolecular NMR spectroscopy. Bioinformatics 31, 1325–1327, (2015).

44 Ziemba, B. P., Booth, J. C. & Jones, D. N. 1H, 13C and 15N NMR assignments of the C1A and C1B subdomains of PKC-delta. Biomol NMR Assign 5, 125–129, (2011).

45 Iwahara, J., Schwieters, C. D. & Clore, G. M. Ensemble approach for NMR structure refinement against (1)H paramagnetic relaxation enhancement data arising from a flexible paramagnetic group attached to a macromolecule. J. Am. Chem. Soc. 126, 5879–5896, (2004).

46 Hatzakis, E., Agiomyrgianaki, A., Kostidis, S. & Dais, P. High-Resolution NMR Spectroscopy: An Alternative Fast Tool for Qualitative and Quantitative Analysis of Diacylglycerol (DAG) Oil. J. Am. Oil Chem. Soc. 88, 1695–1708, (2011).

47 PROTEUM - Bruker AXS (2018). PROTEUM3, Version 2018.1, Bruker AXS Inc., Madison, Wisconsin, USA.

48 Kabsch, W. Xds. Acta Crystallogr D Biol Crystallogr 66, 125–132, (2010).

49 Evans, P. Scaling and assessment of data quality. Acta Crystallogr D Biol Crystallogr 62, 72–82, (2006).

50 Evans, P. R. & Murshudov, G. N. How good are my data and what is the resolution? Acta Crystallogr D Biol Crystallogr 69, 1204–1214, (2013).

51 Emsley, P. & Cowtan, K. Coot: model-building tools for molecular graphics. Acta Crystallogr D Biol Crystallogr 60, 2126–2132, (2004).

52 Liebschner, D. et al. Macromolecular structure determination using X-rays, neutrons and electrons: recent developments in Phenix. Acta Crystallogr D Struct Biol 75, 861–877, (2019).

53 Liebschner, D. et al. Polder maps: improving OMIT maps by excluding bulk solvent. Acta Crystallographica Section D 73, 148–157, (2017).

54 Lebedev, A. A. et al. JLigand: a graphical tool for the CCP4 template-restraint library. Acta Crystallogr D Biol Crystallogr 68, 431–440, (2012).

55 Moriarty, N. W., Grosse-Kunstleve, R. W. & Adams, P. D. electronic Ligand Builder and Optimization Workbench (eLBOW): a tool for ligand coordinate and restraint generation. Acta Crystallogr D Biol Crystallogr 65, 1074–1080, (2009).

56 Pettersen, E. F. et al. UCSF Chimera—A visualization system for exploratory research and analysis. Journal of Computational Chemistry 25, 1605–1612, (2004).

57 Molecular Operating Environment (MOE), 2019.01; Chemical Computing Group ULC, 1010 Sherbooke St. West, Suite #910, Montreal, QC, Canada, H3A 2R7, 2021.

58 Laskowski, R. A. & Swindells, M. B. LigPlot+: Multiple Ligand–Protein Interaction Diagrams for Drug Discovery. Journal of Chemical Information and Modeling 51, 2778–2786, (2011).

