## Supplementary information for "Structural anatomy of C1 domain interactions with DAG and other agonists"

**This file contains:**

Referenced extended discussion (pages 2-7) and crystallography statistics tables (pages 8-9).

**Extended discussion**

**1. C1 consensus sequence and the role of individual C1 residues**

In addition to the two $Zn^{2+}$-coordinating $Cys_3His$ motifs, DAG-sensitive C1 domains share a well-defined consensus amino acid sequence[1]. This consensus signature suggests that sidechain-specific interactions are essential for binding the membrane-embedded ligands. Yet, with the notable exception of DAG interactions with the Gln257 sidechain in the "*sn*-1" mode C1B-DAG complex, none of the ligands engage in sidechain-specific hydrogen bonds with protein residues. All H-bonding interactions are mediated exclusively by the polar backbone atoms of residues Thr242, Leu251, and Gly253. Our structures of multiple C1Bδ-ligand complexes now reveal the specific roles of conserved residues in those interactions for the first time. In doing so, this information not only provides a functional rationale for the consensus sequence of DAG-sensitive C1 domains, but it also explains the outcomes of extensive C1/PKC mutagenesis experiments conducted by multiple groups[2-6].

The four strictly conserved non-$Zn^{2+}$ coordinating residues are Pro241, Gly253, and the Gln257-Gly258 forming the "QG" motif (**Ext. Fig. 9f**, red). Pro241 anchors the carbonyl-containing moieties of DAG and exogenous agonists to the β12 loop (**Ext. Fig. 4c, 9b-e**). In addition to stereospecific H-bonding interactions with the ligands (*vide supra*), Gly253 forms the "depression" in the loop β34 surface that accommodates the hydrophobic moieties of DAG (**Fig. 2b,c**), of the exogenous ligands (**Ext. Fig. 8c**), and potentially of the lipids (**Fig. 4c**) to create a continuous hydrophobic surface. Gln257 stabilizes the loop region through the formation of inter-loop hydrogen bonds and, in the case of DAG, makes a hydrogen bond with the ligand (**Fig. 2b** and **Ext. Fig. 2**). The subsequent glycine residue, Gly258, ensures the conformational flexibility of the β34 loop hinge[7,8] that likely facilitates the membrane recruitment.

Of the nine conserved hydrophobic residues, six form the hydrophobic protein core (**Ext. Fig. 9f**) in the structural $Zn^{2+}$ region (Phe233, Met266, Val268, and Val276) and the inter-loop (Tyr238 and Leu251) regions. The sidechains of the other three (Leu250, Trp252, and Leu254), together with strictly conserved Pro241 and the consensus aromatic residue Phe243, form the outside hydrophobic "cage" that envelopes the bound ligands (**Fig. 9g,h**). One interesting feature of the "cage" is that, along with the expected inter-residue hydrophobic contacts, there is an opportunity for Leu250 to engage in CH-π interactions[9] with the aromatic rings of Trp252 and Phe243. Overall, the spatial organization of these five residues enables the loop region of C1B to effectively interface with lipids, while "shielding" the hydrophilic ligand moieties from the hydrophobic membrane environment (**Fig. 2b,c**; **Fig. 4**). Of note, in the complex of C1Bδ with a water-soluble hydrophilic phorbol ester crystallized in the absence of lipids/detergents, the Trp252 sidechain is oriented away from the membrane-binding loop region (**Fig. 9i**), as is observed in apo C1Bδ (**Fig. 2a**). The support for Trp252 reorientation upon formation of the ternary C1-ligand-membrane complex in solution comes from the unusually large upfield chemical shift change experienced by the Leu250 methyl groups (**Fig. 2e, inset**). This shift is consistent with the aromatic ring current effect, induced by the Trp252 sidechain being brought into close proximity to the Leu250 methyl groups upon DAG binding.

**2. C1 domain interactions with lipids: implications for differential DAG affinities of PKC isoforms.**

The packing arrangements of the C1Bδ chains in the crystal, and their amphiphilic surfaces, create opportunistic binding sites for external DAG and membrane-mimicking agents. There are two peripheral DAG molecules anchored to the groove formed between C1Bδ chains 2 and 3 via hydrogen bonds and nonpolar contacts (**Ext. Fig. 5**). The hydrogen-bonding interactions are mediated by water molecules that bridge DAG oxygen atoms and the polar backbone atoms of Leu250, Leu254, and Val255. The sidechains of these three residues, along with those of Phe243 and Trp252, engage in nonpolar interactions with DAG acyl chains. In addition to peripheral DAG, six DPC molecules engage in non-polar and water-mediated polar contacts with the C1Bδ loop region surface (**Ext. Fig. 6**).

In the C1Bδ-exogenous agonist complexes, we observed peripheral association of DHPC with protein (**Ext. Fig. 10**). DHPC molecules are observed either on the outside of loop β12 (I3A complex), loop β34

(Prostratin and AJH-836 model 7LF3), or both loops (PDBu and ligand-free chain of the AJH-836 model 7LEO) (**Ext. Fig. 10a-e**). The DHPC-protein interactions involve hydrogen bonds, both direct and water-mediated, as well as hydrophobic contacts. The structure of the AJH-836, where one protein chain has a bound ligand and the other ligand-free chain interacts peripherally with DHPC, highlights the versatility of Trp252-lipid interactions (**Ext. Fig. 10f**).

The Trp lipophilic properties are directly relevant to the question of DAG sensitivity of PKC isoforms. Indeed, the identity of an aromatic residue at position 252 or equivalent has a profound effect on the C1B DAG affinity and hence the DAG sensitivity of the parent PKC[5,7,8,10,11]. The novel ($Ca^{2+}$-independent) PKC isoforms have a Trp at that position and show high DAG affinities, while the C1B of conventional ($Ca^{2+}$-dependent) isoforms have a Tyr and show lower DAG affinities. It is suggested that the higher DAG affinities of novel PKC isoform C1B domains compensate for the lack of $Ca^{2+}$-dependent C2 domains that co-target conventional PKCs to anionic membranes[11].

The interaction pattern of Trp252 with ligands (**Fig. 2b,c; Fig. 3a; Fig. 4b,c**) and its membrane insertion (**Fig. 3b**) now makes it possible to rationalize why having Trp over Tyr is thermodynamically advantageous for C1B domains. Compared to the Tyr sidechain, Trp has higher hydrophobicity[12], aromaticity[13,14], and larger electric dipole moment[15]. The hydrophobicity factor is relevant in membrane partitioning[5], and in engaging both "cage" residues (**Ext. Fig. 9g**) and ligands (**Ext. Fig. 4a,b; Ext. Fig. 8b,c**). The extended aromatic system with two fused rings enables cation-π[13] and CH-π[9] interactions that are observed even in the crystalline state in the absence of a fully formed membrane environment (**Ext. Fig. 9g; Ext. Fig. 10f**). In addition, the ~two-fold larger electric dipole moment of the Trp sidechain likely facilitates charge-dipole and dipole-dipole interactions in the complex electrostatic environment of the membrane headgroup region and its interfacial water molecules.

#### 3. C1-DAG lactone interactions

The development of DAG lactones for the modulation of PKC activity was pioneered by the Blumberg and Marquez laboratories[16-25]. The concept behind the development of this class of compounds is that, compared to the endogenous ligand DAG, the rigid cyclic lactone structure reduces the entropic penalty for binding lactones to C1 domains. Furthermore, the hydrophobicity of DAG lactones can be varied via *sn-1/2* linked substituents R1 and R2. Combinatorial library built upon the lactone template identified the R1/R2 substituent combinations that produced distinct and specific cellular responses, attesting to the promise of this class of compounds as modulators of DAG effector proteins[26]. The lactone AJH-836[27] used for our structural studies has one of the highest reported affinities among this class of compounds, and AJH-836 shows selectivity towards the novel PKC isozymes δ and ε relative to conventional α and βII[16].

DAG lactones can bind to the C1 domains in two orientations: (i) "*sn-1*", where the carbonyl of the *sn*-1 ester group hydrogen-bonds with the amide N-H group of Gly253, and (ii) "*sn-2*", where the carbonyl oxygen of the lactone ring hydrogen-bonds with the N-H of Gly253. Previous docking studies, conducted using the Phorbol 13-monoacetate complexed C1Bδ structure[28] suggested that the "*sn*-2" binding mode is preferred for the DAG lactones[29]. Notably, the Trp252 sidechain is oriented away from the membrane-binding loops in that C1Bδ structure (**Ex. Fig. 9i**). Our structure of the C1Bδ-AJH-836 complex, where the Trp252 sidechain is oriented towards the membrane-binding region, has the ligand bound in the "*sn*-1" mode (**Fig. 4**). The preference for the "*sn*-1" mode likely originates from the favorable positioning of bulky R1 and R2 groups relative to the hydrophobic sidechains of the rim residues. Specifically, R1 is sandwiched between Leu254 and Trp252, and R2 contacts Met239, Leu254, and Val255. This arrangement maximizes the protein-ligand hydrophobic contacts and creates a contiguous lipophilic surface for membrane interactions. In this context, the significance of the Trp252 sidechain reorientation (**Fig. 2a**) upon complexation with PKC agonists becomes clear. When oriented towards the loop region, Trp252 displays considerable functional versatility as it can interface simultaneously with the hydrophobic ligand and the surrounding lipids (*vide supra*).

### References


1 Hurley, J. H., Newton, A. C., Parker, P. J., Blumberg, P. M. & Nishizuka, Y. Taxonomy and function of C1 protein kinase C homology domains. *Protein Sci.* **6**, 477-480, (1997).

2 Kazanietz, M. G. *et al.* Residues in the second cysteine-rich region of protein kinase C delta relevant to phorbol ester binding as revealed by site-directed mutagenesis. *J. Biol. Chem.* **270**, 21852-21859, (1995).

3 Choi, Y. *et al.* Conformationally Constrained Analogues of Diacylglycerol (DAG). 28. DAG-dioxolanones Reveal a New Additional Interaction Site in the C1b Domain of PKCδ. *J. Med. Chem.* **50**, 3465-3481, (2007).

4 Rahman, G. M. *et al.* Identification of the activator-binding residues in the second cysteine-rich regulatory domain of protein kinase Ctheta (PKCtheta). *Biochem. J.* **451**, 33-44, (2013).

5 Stewart, M. D., Cole, T. R. & Igumenova, T. I. Interfacial partitioning of a loop hinge residue contributes to diacylglycerol affinity of conserved region 1 domains. *J. Biol. Chem.* **289**, 27653-27664, (2014).

6 Wang, Q. J. *et al.* Role of hydrophobic residues in the C1b domain of protein kinase C delta on ligand and phospholipid interactions. *J Biol Chem* **276**, 19580-19587, (2001).

7 Stewart, M. D., Morgan, B., Massi, F. & Igumenova, T. I. Probing the determinants of diacylglycerol binding affinity in the C1B domain of protein kinase Calpha. *J. Mol. Biol.* **408**, 949-970, (2011).

8 Stewart, M. D. & Igumenova, T. I. Toggling of Diacylglycerol Affinity Correlates with Conformational Plasticity in C1 Domains. *Biochemistry* **56**, 2637-2640, (2017).

9 Plevin, M. J., Bryce, D. L. & Boisbouvier, J. Direct detection of CH/pi interactions in proteins. *Nat Chem* **2**, 466-471, (2010).

10 Dries, D. R., Gallegos, L. L. & Newton, A. C. A single residue in the C1 domain sensitizes novel protein kinase C isoforms to cellular diacylglycerol production. *J. Biol. Chem.* **282**, 826-830, (2007).

11 Giorgione, J. R., Lin, J. H., McCammon, J. A. & Newton, A. C. Increased membrane affinity of the C1 domain of protein kinase Cdelta compensates for the lack of involvement of its C2 domain in membrane recruitment. *J Biol Chem* **281**, 1660-1669, (2006).

12 Wimley, W. C. & White, S. H. Experimentally determined hydrophobicity scale for proteins at membrane interfaces. *Nat Struct Biol* **3**, 842-848, (1996).

13 Dougherty, D. A. Cation-pi interactions involving aromatic amino acids. *J Nutr* **137**, 1504S-1508S; discussion 1516S-1517S, (2007).

14 de Jesus, A. J. & Allen, T. W. The role of tryptophan side chains in membrane protein anchoring and hydrophobic mismatch. *Biochim Biophys Acta* **1828**, 864-876, (2013).

15 Yau, W. M., Wimley, W. C., Gawrisch, K. & White, S. H. The preference of tryptophan for membrane interfaces. *Biochemistry* **37**, 14713-14718, (1998).

16 Cooke, M. *et al.* Characterization of AJH-836, a diacylglycerol-lactone with selectivity for novel PKC isozymes. *J. Biol. Chem.* **293**, 8330-8341, (2018).

17 Ohashi, N. *et al.* Synthesis and Evaluation of Dimeric Derivatives of Diacylglycerol-Lactones as Protein Kinase C Ligands. *Bioconjug Chem* **28**, 2135-2144, (2017).

18 Nomura, W. *et al.* Synthetic caged DAG-lactones for photochemically controlled activation of protein kinase C. *Chembiochem* **12**, 535-539, (2011).

19 Malolanarasimhan, K. *et al.* Conformationally constrained analogues of diacylglycerol (DAG). 27. Modulation of membrane translocation of protein kinase C (PKC) isozymes alpha and delta by diacylglycerol lactones (DAG-lactones) containing rigid-rod acyl groups. *J Med Chem* **50**, 962-978, (2007).

20 Pu, Y. *et al.* A novel diacylglycerol-lactone shows marked selectivity in vitro among C1 domains of protein kinase C (PKC) isoforms alpha and delta as well as selectivity for RasGRP compared with PKCalpha. *J Biol Chem* **280**, 27329-27338, (2005).

21 Lee, J. *et al.* Conformationally constrained diacylglycerol (DAG) analogs: 4-C-hydroxyethyl-5-O-acyl-2,3-dideoxy-D-glyceropentono-1,4-lactone analogs as protein kinase C (PKC) ligands. *Eur J Med Chem* **39**, 69-77, (2004).
22 Choi, Y. *et al.* Conformationally constrained analogues of diacylglycerol. 19. Synthesis and protein kinase C binding affinity of diacylglycerol lactones bearing an N-hydroxylamide side chain. *J Med Chem* **46**, 2790-2793, (2003).
23 Nacro, K., Bienfait, B., Lewin, N. E., Blumberg, P. M. & Marquez, V. E. Diacylglycerols with lipophilically equivalent branched acyl chains display high affinity for protein kinase C (PK-C). A direct measure of the effect of constraining the glycerol backbone in DAG lactones. *Bioorg Med Chem Lett* **10**, 653-655, (2000).
24 Lee, J. *et al.* Conformationally constrained analogues of diacylglycerol. 12. Ultrapotent protein kinase C ligands based on a chiral 4,4-disubstituted heptono-1,4-lactone template. *J Med Chem* **39**, 36-45, (1996).
25 Lee, J., Marquez, V. E., Blumberg, P. M., Krausz, K. W. & Kazanietz, M. G. Conformationally constrained analogues of diacylglycerol (DAG)--II. Differential interaction of delta-lactones and gamma-lactones with protein kinase C (PK-C). *Bioorg Med Chem* **1**, 119-123, (1993).
26 Duan, D. *et al.* Conformationally constrained analogues of diacylglycerol. 29. Cells sort diacylglycerol-lactone chemical zip codes to produce diverse and selective biological activities. *J Med Chem* **51**, 5198-5220, (2008).
27 Ann, J. *et al.* Design and synthesis of protein kinase C epsilon selective diacylglycerol lactones (DAG-lactones). *Eur J Med Chem* **90**, 332-341, (2015).
28 Zhang, G. G., Kazanietz, M. G., Blumberg, P. M. & Hurley, J. H. Crystal-Structure of the Cys2 Activator-Binding Domain of Protein-Kinase C-Delta in Complex with Phorbol Ester. *Cell* **81**, 917-924, (1995).
29 Sigano, D. M. *et al.* Differential binding modes of diacylglycerol (DAG) and DAG lactones to protein kinase C (PK-C). *J. Med. Chem.* **46**, 1571-1579, (2003).

**Supporting information Table 1 |** Crystallography data collection, and refinement statistics of apo C1Bδ and its complexes with diacylglycerol and AJH-836.

| | **Apo (7KND)** | **Diacylglycerol (7L92)** | **AJH 836-DHPC (7LEO)** | **AJH 836 (7LF3)** |
|---|---|---|---|---|
| **Data Collection** | | | | |
| Space group | P 41 | H 3 | C 1 2 1 | C 1 2 1 |
| Cell dimensions | | | | |
| *a*, *b*, *c* (Å) | 37.26, 37.26, 31.89 | 89.07, 89.07, 218.68 | 83.39, 50.89, 37.48 | 34.91, 25.94, 57.64 |
| *α*, *β*, *γ* (°) | 90.00, 90.00, 90.00 | 90.00, 90.00, 120.00 | 90.00, 107.7 90.00 | 90.00, 92.57, 90.00 |
| Resolution (Å) | 37.26–1.39 (1.41-1.39) | 44.60 – 1.75 (1.78-1.75) | 50 – 1.65 (1.68 – 1.65) | 28.79 – 1.13 (1.15 – 1.13) |
| $R_{\text{merge}}$ | 0.056 (1.65) | 0.07 (0.613) | 0.1 (0.831) | 0.13 (2.3) |
| $I/\sigma(I)$ | 15 (0.8) | 6.8 (1.6) | 7.4 (1.77) | 15.5 (0.6) |
| Completeness (%) | 100 (100) | 99.1 (99.6) | 98.5 (98.1) | 95.6 (86.6) |
| Redundancy | 6.3 (4.3) | 2.8 (2.8) | 5.4 (3.7) | 5.7 (5.5) |
| CC1/2 | 0.995 (0.522) | | 0.99 (0.568) | 0.98 (0.396) |
| **Refinement** | | | | |
| Resolution (Å) | 37.26 – 1.39 | 28.9-1.75 | 29.96 – 1.65 | 28.79 – 1.13 |
| No. reflections | 8897 | 64618 | 17697 | 18562 |
| $R_{\text{work}}$ / $R_{\text{free}}$ | 0.216 / 0.246 | 0.2142 / 0.2460 | 0.2237 / 0.2439 | 0.1826 /0.1993 |
| No. atoms | | | | |
| Protein | 422 | 3371 | 819 | 440 |
| Solvent | 26 | 231 | 61 | 51 |
| Ligands/metals | 2 | 400 | 95 | 62 |
| *B* factors (Å$^2$) | | | | |
| Protein | 31 | 36 | 33 | 18 |
| Ligands/metals | 25 | 39 | 38.3 | 31.5 |
| R.m.s. deviations | | | | |
| Bond lengths (Å) | 0.01 | 0.009 | 0.006 | 0.006 |
| Bond angles (°) | 1.35 | 1.27 | 1.1 | 0.942 |

**Supporting information Table 2 |** Crystallography data collection, and refinement statistics of the C1Bδ complexes with PDBu, Prostratin, and Ingenol-3-angelate.

| | **Phorbol 12,13-dibutyrate (7KNJ)** | **Prostratin (7LCB)** | **Ingenol-3-angelate (7KO6)** |
|---|---|---|---|
| **Data Collection** | | | |
| Space group | C 1 2 1 | C 1 2 1 | C 1 2 1 |
| Cell dimensions | | | |
| $a$, $b$, $c$ (Å) | 34.20, 26.14, 58.16 | 34.40, 25.87, 57.96 | 34.38, 25.91, 57.73 |
| $\alpha$, $\beta$, $\gamma$ (°) | 90.00, 96.92, 90.00 | 90.00, 97.29, 90.00 | 90.00, 98.39, 90.00 |
| Resolution (Å) | 57.73–1.57 (1.67-1.57) | 57.50 – 1.70 (1.8 – 1.7) | 57.11 – 1.80 (1.90 - 1.80) |
| $R_{\text{merge}}$ | 0.055 (0.2054) | 0.06 (0.2636) | 0.1144 (0.4800) |
| $I/\sigma(I)$ | 21.05 (3.28) | 15.48 (3.00) | 15.76 (3.79) |
| Completeness (%) | 85.1 (34) | 89.1 (48.4) | 100 (100.0) |
| Redundancy | 5.71 (0.44) | 3.85 (0.63) | 12.54 (8.97) |
| CC1/2 | | | |
| **Refinement** | | | |
| Resolution (Å) | 19.24-1.57 | 28.75 - 1.7 | 28.56-1.8 |
| No. reflections | 6194 | 5113 | 4815 |
| $R_{\text{work}}$ / $R_{\text{free}}$ | 0.1772 / 0.2236 | 0.1719 / 0.2136 | 0.1757 / 0.2154 |
| No. atoms | | | |
| Protein | 431 | 431 | 423 |
| Solvent | 44 | 38 | 53 |
| Ligands/metals | 97 | 62 | 65 |
| $B$ factors (Å$^2$) | | | |
| Protein | 13 | 13 | 14 |
| Ligands/metals | 23.6 | 20 | 20 |
| R.m.s. deviations | | | |
| Bond lengths (Å) | 0.008 | 0.009 | 0.01 |
| Bond angles (°) | 2.42 | 1.445 | 2.19 |
